# Fine-Tuning α-Synuclein Phase Separation through Sequence-Optimized Peptide Modulators

**DOI:** 10.64898/2026.02.21.707152

**Authors:** Tatsuya Ikenoue, Tsuyoshi Konuma, Takahisa Ikegami, Hiroaki Suga

**Author notes:** These authors contributed equally to this work (Tatsuya Ikenoue, Tsuyoshi Konuma).

## Abstract

Liquid–liquid phase separation (LLPS) of intrinsically disordered proteins underlies the formation of biomolecular condensates that regulate diverse cellular processes, while its dysregulation contributes to protein aggregation and disease. Despite its importance, molecularly defined and target-specific strategies to control LLPS remain limited. Here, we present a systematic framework for designing de novo peptides that induce and modulate LLPS of α-synuclein. By integrating deep mutational scanning with peptide screening, we identified sequence features that govern condensate formation and enabled the creation of optimized peptides with high efficiency and specificity. Biophysical analyses revealed that LLPS efficiency is dictated by the interplay of solubility, multivalency, and cooperative interactions, resulting in a distinctive bell-shaped phase diagram. Thermodynamic measurements and imaging-based analyses further demonstrated that condensate stability and material properties can be rationally tuned through peptide optimization. Together, these findings establish generalizable design principles for engineering LLPS modulators in biologically and pathologically relevant protein systems.

## Introduction

Liquid–liquid phase separation (LLPS) of intrinsically disordered proteins has emerged as a fundamental organizing principle of biomolecular condensates^1–4^. These dynamic, membraneless compartments regulate diverse cellular processes by concentrating proteins and nucleic acids into liquid-like droplets, yet dysregulated LLPS is implicated in the pathological aggregation of amyloidogenic proteins, including α-synuclein (αSyn), a hallmark of Parkinson’s disease^5–9^.

At the molecular level, LLPS is driven by multivalent weak interactions among intrinsically disordered regions, including π–π interactions between aromatic residues, cation–π interactions, electrostatic contacts between oppositely charged residues, and dipole–dipole contacts^10–15^. The balance and distribution of these interactions dictate not only whether phase separation occurs but also the physical properties of condensates, such as fluidity, stability, and susceptibility to undergo liquid-to-solid transitions^16–18^. However, because these forces are individually weak and often highly dynamic, it has been challenging to define the sequence determinants that govern LLPS efficiency and selectivity.

Despite the growing recognition of LLPS as central to cellular organization and disease, strategies for its molecular control remain limited^2,19^. Existing modulators—including small molecules^12,20,21^, simplified model macromolecules^22–24^, or polypeptides inspired by native proteins^25–29^—typically act nonspecifically, perturbing physicochemical environments rather than engaging defined molecular targets. Consequently, they offer little capacity to disentangle LLPS from aggregation or to achieve precise, target-specific modulation of condensate behavior. This lack of selectivity is a major bottleneck for basic studies and for therapeutic efforts aimed at pathological condensates^30–35^.

We previously identified de novo peptides capable of inducing LLPS of αSyn^36^. Among them, FL2 displayed modest activity with low specificity, whereas FD1 exhibited strong target selectivity. These peptides established that LLPS can be artificially induced—even in proteins that do not spontaneously phase separate—providing the proof-of-concept for target-specific condensate control. However, the molecular determinants linking peptide sequence to LLPS efficiency and selectivity had not been systematically defined.

Here, we extend this strategy by integrating deep mutational scanning (DMS) with our peptide screening platform^37,38^ to comprehensively analyze and optimize LLPS-inducing peptides. By coupling sequence–activity mapping with detailed biophysical characterization, we uncover principles by which binding affinity, solubility, and multivalency converge to regulate condensate formation. This approach not only enables the creation of optimized peptides with high efficiency and specificity but also provides mechanistic insight into how defined molecular interactions drive or suppress phase separation. More broadly, our study lays the foundation for engineering next-generation LLPS modulators, with implications for mechanistic dissection of condensates and for the development of therapeutic strategies targeting pathological phase transitions^30,31^.

## Results

### Evaluation of LLPS induction under physiological conditions and optimization of peptide scaffold design

We previously identified two representative de novo peptides, FL2 and FD1, with distinct specificities toward α-synuclein (αSyn). FL2 displayed relatively low target selectivity, whereas FD1 exhibited high target specificity in inducing LLPS. In our initial studies, phase separation assays were performed in 20 mM sodium phosphate (NaPi) buffer, which provided robust LLPS induction. However, when the buffer was replaced with phosphate-buffered saline (PBS; 10 mM NaPi, 140 mM NaCl) to better approximate physiological ionic conditions, both peptides exhibited markedly reduced LLPS-inducing efficiency (**Supplementary Fig. 1**). These observations underscore the importance of peptide optimization to achieve robust condensate formation under physiologically relevant conditions. To determine the optimal structural form of peptides for systematic mutational analysis, we next compared cyclic and linear versions of the LLPS-inducing sequences. Using differential interference contrast (DIC) microscopy and turbidimetry, we quantified LLPS efficiency across these two peptide formats (**Supplementary Fig. 2**). The linear peptides (FL2L and FD1L) consistently exhibited enhanced LLPS-inducing activity relative to their cyclic counterparts. Furthermore, constructing tandem-repeat versions of the linear peptides dramatically increased their LLPS efficiency, suggesting that multivalency achieved through linear extension strongly promotes condensate formation. Based on these results, we chose to perform deep mutational scanning (DMS) in the linear peptide format to maximize sensitivity for identifying sequence determinants of LLPS induction.

### Comprehensive analysis of αSyn-binding determinants by deep mutational scanning

To comprehensively map mutation–affinity relationships of LLPS-inducing peptides, we applied a DMS approach based on our previously established in vitro translation–coupled peptide screening system (**Fig. 1a**). Libraries of systematically mutated peptide variants derived from FL2L and FD1L were generated and subjected to screening in the presence of αSyn. The relative enrichment or depletion of each variant before and after selection was quantified by next-generation sequencing, thereby enabling a high-throughput, quantitative assessment of the contribution of individual amino acid substitutions to αSyn binding (**Fig. 1b**).

**Fig. 1.**
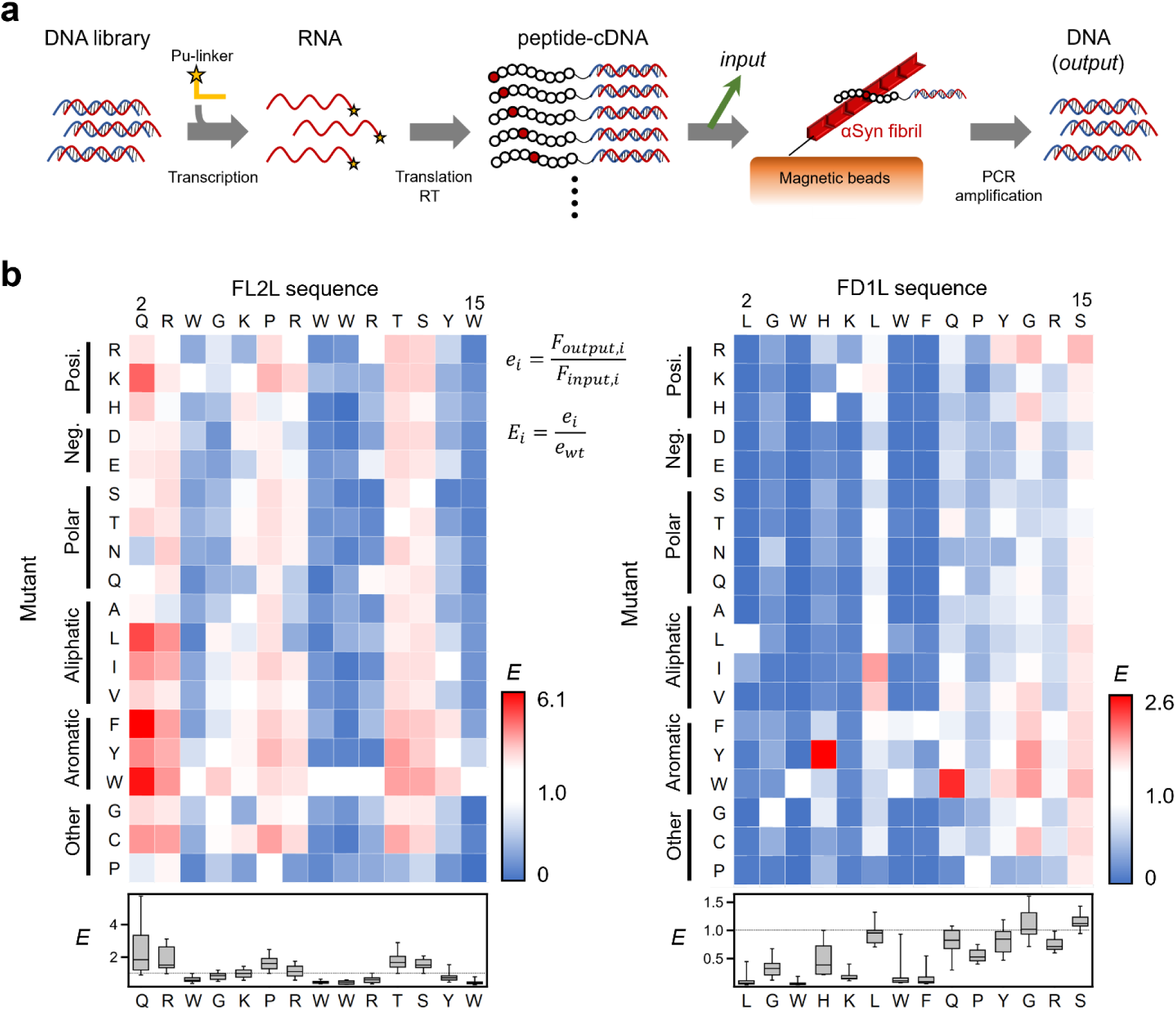
Deep mutational scanning reveals sequence determinants of αSyn-binding LLPS-inducing peptides. **a** Schematic overview of the deep mutational scanning platform developed to systematically evaluate the impact of single-point mutations on peptide affinity for αSyn. Two peptide scaffolds (FL2L and FD1L) were used to construct mRNA display libraries in which each codon was individually randomized using the NNK scheme. The mRNA was ligated to a puromycin-containing linker, translated in vitro, and reverse-transcribed to generate mRNA–peptide–cDNA fusion complexes. These libraries were incubated with αSyn fibrils immobilized on magnetic beads under stringent conditions to isolate binding-competent variants. Bound cDNA species were PCR-amplified, and both input and output libraries were subjected to next-generation sequencing. For each variant (*i*), enrichment scores (*e*) were calculated by comparing the relative abundance (*F*) in the output versus the input, thereby quantifying the impact of each mutation on αSyn binding. **b** Enrichment heatmaps show the positional and amino acid-specific effects on binding affinity (*E*) across the primary sequence of FL2L (left) and FD1L (right). Box-and-whisker plots of *E* scores at each peptide residue are also shown. Mutations that enhance (red) or impair (blue) αSyn interaction were detected, revealing critical contact residues and non-essential regions. These data guided subsequent rational peptide optimization for enhanced LLPS efficiency and solubility.

This DMS analysis revealed distinct sequence features that either enhanced or impaired peptide affinity toward αSyn. Mutations that increased binding affinity highlighted specific “hot spot” residues critical for condensate-promoting interactions, whereas substitutions that consistently reduced affinity delineated peptide regions indispensable for target engagement. Conversely, certain positions tolerated a wide variety of substitutions without appreciable effects on affinity, suggesting that these residues are less critical for αSyn recognition.

Overall, the mutational landscapes of both FL2L and FD1L revealed a convergent trend in which aromatic residues and positively charged residues contributed positively to binding affinity, in agreement with the role of π–π and electrostatic interactions in condensate stabilization (**Supplementary Fig. 3**). Importantly, FD1L showed a narrower mutational tolerance profile than FL2L, consistent with its higher binding specificity. This feature later proved to be critical for developing LLPS-inducing peptides with minimal off-target effects. Together, these findings establish the foundation for sequence-guided peptide optimization and lay the groundwork for dissecting the interplay between αSyn affinity and LLPS-inducing capacity.

### Optimization of LLPS-inducing peptides reveals correlation between affinity, solubility, and condensate properties

Guided by the mutational landscapes obtained from the DMS data set, we selected a panel of 75 peptide variants for chemical synthesis. These included peptides carrying affinity-enhancing substitutions, mutations introduced at positions predicted to be non-essential for binding but potentially relevant to LLPS, and variants lacking solubility tags. The LLPS-inducing efficiencies of these peptides were systematically assessed by DIC microscopy and turbidimetry (**Supplementary Fig. 4**).

Strikingly, in the target-specific FD1 background, peptide affinity toward αSyn correlated strongly with LLPS-inducing capacity (**Fig. 2a**). In addition, peptide solubility emerged as a robust predictor of LLPS efficiency: peptides with lower solubility consistently promoted the formation of condensates more efficiently (**Fig. 2b**). However, the excessively reduced solubility led to the formation of gel-like assemblies or insoluble aggregates rather than liquid-like droplets (**Fig. 2c and Supplementary Fig. 5**). Notably, in the FL2L scaffold, substitutions involving proline represented a clear exception to this trend. Replacement of proline residues with other amino acids promoted aggregation irrespective of the predicted solubility, indicating that proline plays an exceptional role in suppressing aggregation beyond its contribution to solubility (**Fig. 2b, c**). Peptide solubility scores were further evaluated using the CamSol algorithm^39^, and fluorescence recovery after photobleaching (FRAP) measurements revealed that droplet fluidity was also strongly correlated with peptide solubility (**Fig. 2d, e**).

**Fig. 2.**
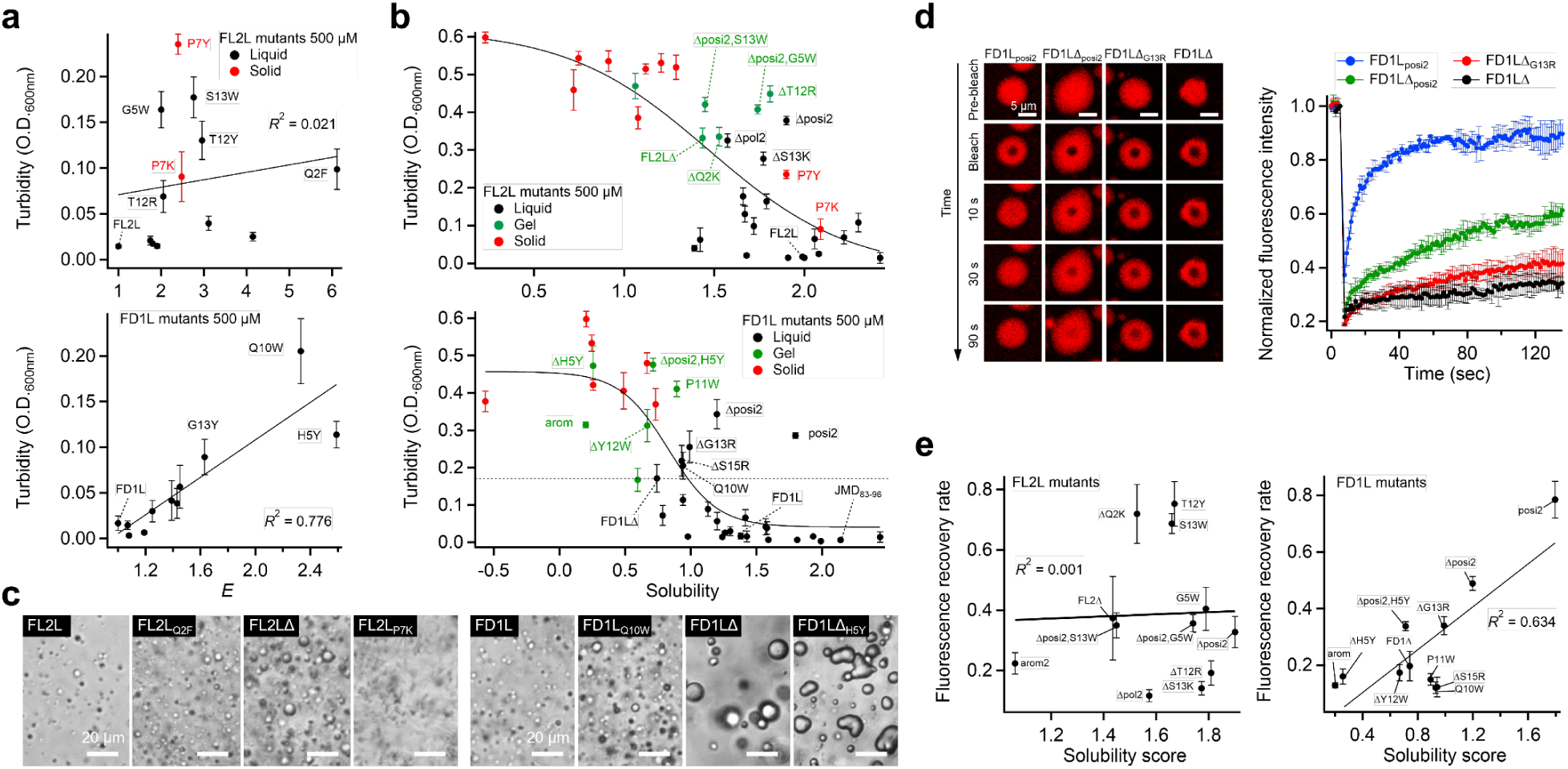
Mutational analysis reveals solubility and sequence determinants governing LLPS efficiency and physical properties of αSyn-peptide condensates. **a** Correlation between enrichment score (*E*) derived from deep mutational scanning and turbidity-based LLPS efficiency across 75 chemically synthesized peptide variants. A positive correlation was particularly evident for FD1L-based peptides, indicating that increased αSyn-binding affinity enhances LLPS induction. **b** Relationship between peptide solubility (as predicted by CamSol scores^39^) and LLPS efficiency. Variants with lower solubility tended to show higher turbidity, reflecting increased LLPS propensity. However, excessively insoluble peptides led to gel-like assemblies or aggregates rather than liquid-like droplets. Color coding indicates droplet types: black (liquid), green (gel), red (aggregate). Solid lines are included for visual guidance. **c** Representative DIC images of αSyn condensates induced by selected peptide variants highlight diverse morphological outcomes. FL2L_Q2F_ and FD1L_Q10W_ represent high-*E* variants; FL2LΔ and FD1LΔ represent solubility-reduced peptides; FL2L_P7K_ and FD1LΔ_H5Y_ display altered condensate morphology. **d** Representative fluorescence images of αSyn-peptide droplets obtained during the FRAP analysis following 1 h incubation at room temperature. The FRAP recovery curves indicate that the liquidity of the droplets varies depending on the peptide variant. **e** Correlation between peptide solubility and fluorescence recovery rate from FRAP, linking solubility to the dynamic properties of condensates.

Through this optimization process, we identified two variants—FL2LΔ_posi2_ and FD1L_posi2_—that exhibited dramatically enhanced LLPS efficiency while maintaining sufficient solubility to selectively promoted the droplet formation without aggregation. Scrambled controls confirmed that the activity of these optimized peptides was sequence-specific, with FD1L_posi2_ in particular displaying high sequence selectivity (**Supplementary Fig. 6**). When benchmarked against model peptides such as poly-lysines (poly-K) and the previously reported LLPS-inducing peptide JMD_83–96_ (derived from VAMP2)^40^, our optimized peptides demonstrated substantially superior LLPS efficiency, highlighting the advantage of sequence-guided optimization over nonspecific or naturally derived motifs.

Collectively, this section establishes that αSyn LLPS efficiency can be dramatically improved through systematic peptide sequence optimization. The results also highlight the interplay between affinity, solubility, and droplet fluidity—providing key design principles for next-generation LLPS modulators with defined physicochemical properties and molecular specificity.

### Optimized peptides reveal bell-shaped phase diagrams and reversible LLPS behavior

Because multivalent interactions are central to LLPS, we hypothesized that increasing the number of repeated peptide motifs would enhance condensate formation. Based on this rationale, seven peptide variants were selected according to their solubility and LLPS-inducing efficiency, and tandem-repeat sequences were designed. Due to the practical limitations of solid-phase peptide synthesis (SPPS), tandem-repeat peptides were synthesized only as two-fold repeats.

These dimeric peptides exhibited a pronounced enhancement in LLPS efficiency, exceeding that expected from a simple twofold increase in peptide concentration (**Fig. 3a, b, Supplementary Fig. 7**). However, tandem repetition also tended to reduce droplet fluidity, suggesting the formation of more tightly networked condensates. Among them, (FD1L_posi2_)_2_ retained high fluidity while maintaining strong LLPS-inducing capacity.

**Fig. 3:**
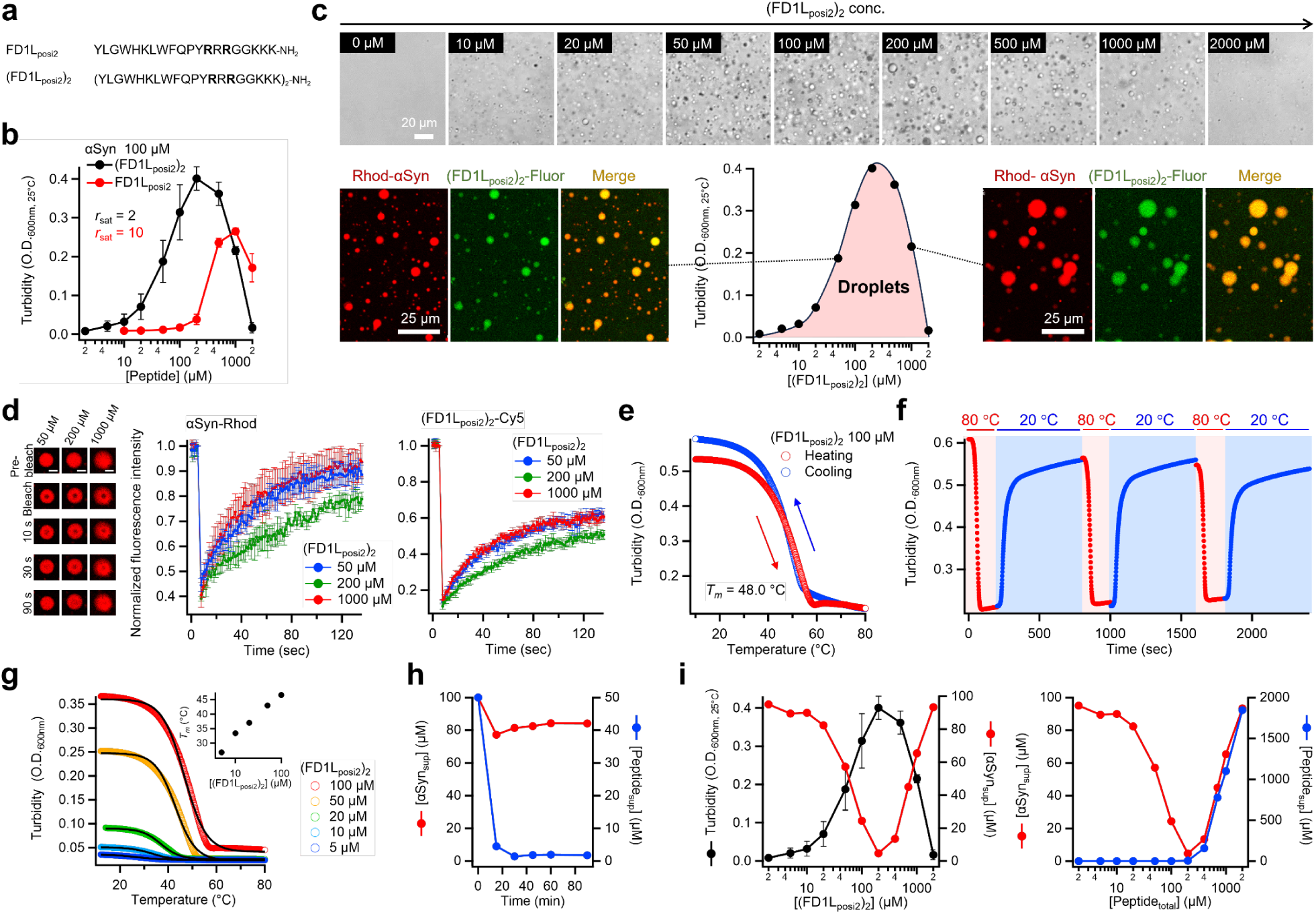
Optimized peptides significantly enhance LLPS efficiency and exhibit bell-shaped phase diagram. **a** Amino acid sequence of the optimized peptide FD1L_posi2_, which enhances both LLPS efficiency and droplet fluidity. **b** Phase diagrams of αSyn LLPS induced by (FD1L_posi2_)_2_ and FD1L_posi2_, illustrating concentration-dependent phase behavior. **c** Representative DIC and fluorescence images showing concentration-dependent changes in droplet formation upon treatment with (FD1L_posi2_)_2_. **d** Representative fluorescence images of αSyn–Rhod obtained during the FRAP experiments, together with FRAP recovery curves of αSyn–Rhod and (FD1L_posi2_)_2_–Cy5. Measurements were performed after 1 h incubation at room temperature at varying concentrations of (FD1L_posi2_)_2_, highlighting the high internal dynamics and fluidity of the droplets. **e** Thermal dissociation and re-condensation of LLPS droplets monitored by turbidity. The temperature scan rate was 5 °C/min. **f** Reversible thermal behavior of the droplets assessed by repeated heating and cooling cycles, demonstrating thermal reversibility. **g** Thermal stability of αSyn droplets induced by various concentrations of (FD1L_posi2_)_2_. Melting temperature (*T*_m_), shown in the inlet, were determined by fitting the turbidity curves (solid lines). **h** Concentrations of αSyn and (FD1L_posi2_)_2_ remaining in the supernatant (αSyn_sup_ and Peptide_sup_, respectively) after centrifugation at 11,000×*g*, determined by HPLC. LLPS was induced using 50 μM (FD1L_posi2_)_2_. **i** Residual concentrations of αSyn and (FD1L_posi2_)_2_ in the supernatant after 30 min incubation with varying peptide concentrations, quantifying LLPS efficiency.

The favorable physicochemical profile of (FD1L_posi2_)_2_ allowed us to investigate condensate behavior at high peptide concentrations without interference from aggregation. When (FD1L_posi2_)_2_ was titrated across concentrations ranging from 2 μM to 2,000 μM in the presence of αSyn, turbidimetry revealed a bell-shaped phase diagram (**Fig. 3c**). Droplet formation increased with peptide concentration until a saturation point was reached, beyond which further peptide addition led to dispersal of αSyn into the dilute phase and a progressive decrease of condensates. At the highest concentration tested (2,000 μM), phase separation was completely abolished, and αSyn remained fully dispersed.

FRAP confirmed that droplets formed under these conditions retained high fluidity (**Fig. 3d**). Moreover, droplet assembly and dissolution were shown to be reversible, as temperature cycling repeatedly induced disassembly and reformation of condensates in a manner reminiscent of protein folding–unfolding transitions (**Fig. 3e, f**). The midpoint of dispersal allowed determination of an apparent melting temperature (*T*_m_), which increased with peptide concentration (**Fig. 3g**).

To further quantify the thermodynamics of phase separation, we measured αSyn concentrations in the supernatant after centrifugation by HPLC (**Fig. 3h, i, Supplementary Fig. 8**). LLPS induction reached apparent equilibrium within 15 min. At the critical peptide concentration, residual αSyn in the dilute phase stabilized at ∼5 μM, a value comparable to the stability of amyloid fibrils. Importantly, while the residual peptide concentration in the dilute phase remained constant (∼2 μM) below the critical threshold, excess peptide accumulated in the dilute phase above this point, consistent with the saturation-driven shift in phase behavior.

The above data highlight how rational peptide optimization enables finely tunable, reversible LLPS with thermodynamically stable droplets. Moreover, the bell-shaped response illustrates a mechanistic transition: from efficient networked co-condensation at stoichiometric balance to dispersive inhibition at excess ligand, thereby providing critical insight into the concentration dependence of LLPS in biomolecular systems.

### Interaction analyses reveal molecular determinants of phase separation

To dissect the molecular mechanism underlying peptide-induced phase separation, we performed isothermal titration calorimetry (ITC) to evaluate thermodynamic parameters and binding stoichiometry (**Fig. 4a**). The reactions encompassing both binding and phase separation progressed with a protein-to-peptide stoichiometry of 2:1. Compared with amyloid fibril formation, the interaction was characterized by a stronger enthalpic contribution, highlighting the importance of favorable intermolecular contacts in promoting condensate assembly^41,42^.

**Fig. 4:**
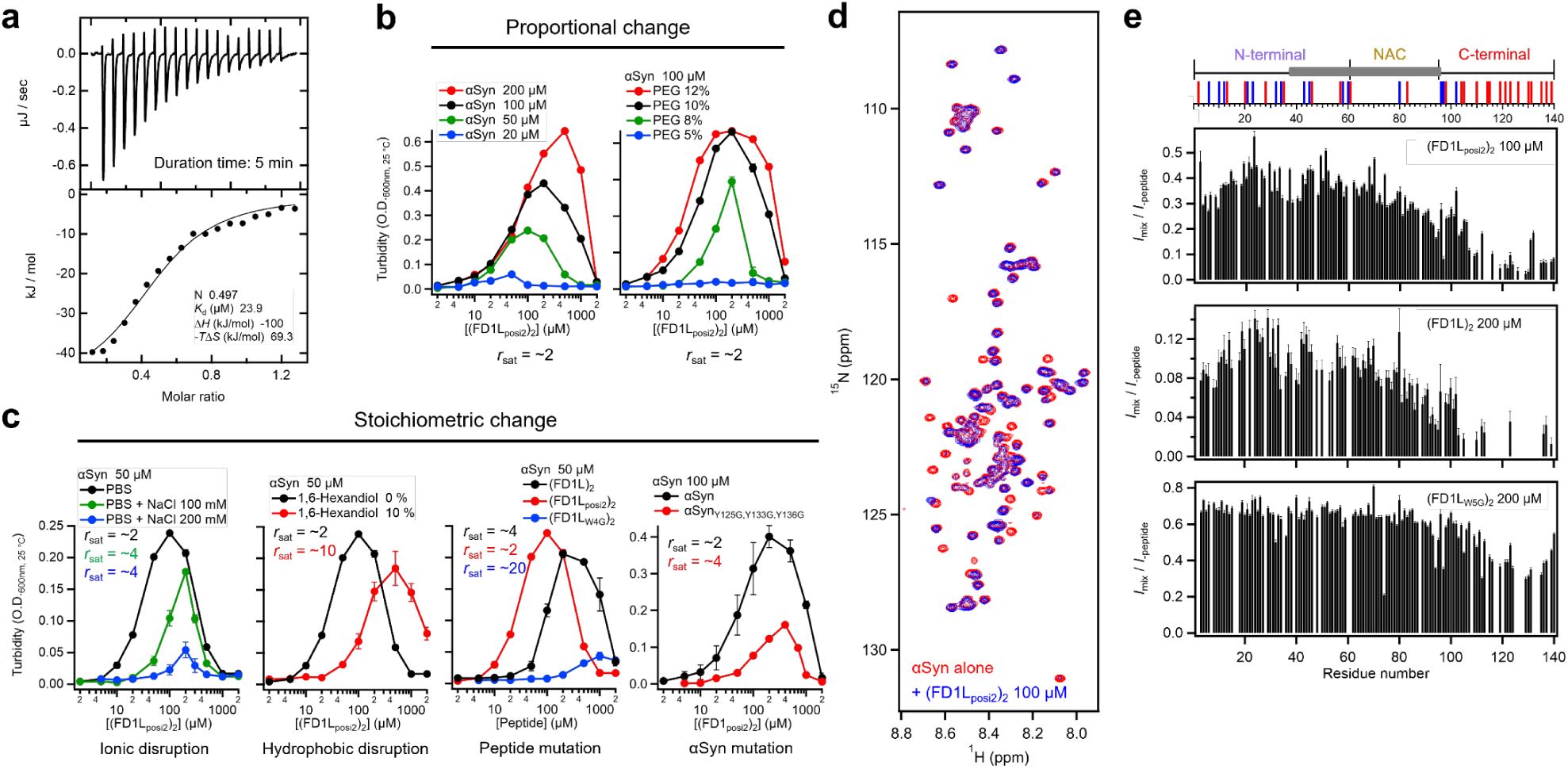
Interaction mode of the optimized peptides. **a** ITC thermogram showing the titration of (FD1L_posi2_)_2_ into αSyn monomers. Peptides at concentrations of 200 μM dissolved in PBS buffer with 1% DMSO were titrated to the 50 μM of αSyn in the sample cell at 25 °C. The data were fitted using an independent one-binding site model. **b** Phase diagrams were compared under varying protein concentrations, PEG concentrations, and in the presence of 1 mM ubiquitin. In all cases, the αSyn:peptide stoichiometry at saturation remained 1:2, indicating that crowding effects or addition of non-interacting proteins do not alter the fundamental binding stoichiometry. **c** Phase boundary analysis under different ionic strengths and in the presence of 1,6-hexanediol revealed shifts of the LLPS critical concentration toward higher peptide concentrations. These perturbations indicate that electrostatic and hydrophobic contributions are essential for condensate stability, and that disruption of these interactions alters the phase diagram. Phase diagrams for the unmodified peptide (FD1L)_2_ and a variant containing a deleterious mutation (FD1L_W4G_)_2_ are also included for comparison. Mutational analysis of αSyn revealed that substitution of Y125, Y133, and Y136 significantly weakened binding reduced LLPS efficiency. **d** ^1^H-^15^N-HSQC spectra of 100 μM ^15^N-labeled αSyn monomers in the absence (red) and presence (blue) of peptides. **e** Peak intensity ratios (*I*_+peptide_*/I*_-peptide_) derived from the HSQC spectra, indicating changes in site-specific interactions upon peptide binding. Distribution of charged residues along the αSyn sequence is shown above the spectra. Negatively and positively charged residues at neutral pH are indicated by red and blue bars, respectively. The fibril core region is highlighted by gray rectangles.

Perturbation experiments confirmed that this stoichiometry was robust as long as the system was under conditions that did not directly affect peptide–protein interactions. Variations in αSyn concentration, PEG concentration, and the presence of ubiquitin or tRNA did not alter the saturation point, which remained fixed at a peptide-to-protein ratio (*r*_sat_) of 2:1 (**Fig. 4b, Supplementary Fig. 9**). In contrast, increasing ionic strength by raising the salt concentration or adding 1,6-hexanediol significantly shifted the critical concentration, moving the phase boundary toward higher peptide concentrations (**Fig. 4c, Supplementary Fig. 10**). Because 1,6-hexanediol is known to weaken hydrophobic interactions, and elevated salt concentrations attenuate long-range electrostatic attractions, these results indicate that both electrostatic and hydrophobic interactions play essential roles in stabilizing peptide-induced condensates. Moreover, peptide mutations that directly disrupted binding substantially reshaped the phase diagram. For example, substitution of the critical residue W4G resulted in a pronounced reduction in LLPS efficiency and shifted the critical concentration to higher values. Consistently, αSyn mutants carrying aromatic-to-glycine substitutions also exhibited a pronounced loss of LLPS efficiency, confirming that these residues are essential for stabilizing peptide-mediated condensates.

Nuclear magnetic resonance (NMR) ^1^H–^15^N heteronuclear single-quantum correlation (HSQC) spectra of αSyn revealed that the optimized peptides interact with the negatively charged C-terminal region of αSyn, similar to the parental (FD1L)_2_ peptide (**Fig. 4d, e, Supplementary Fig. 11a**). Notably, optimization involved substitutions at positions outside the primary interaction interface, indicating that the overall binding mode remained unchanged. Likewise, tandem-repeat variants did not introduce major alterations in interaction patterns. In contrast, the W4G mutation weakened binding, emphasizing the critical role of aromatic residues in maintaining multivalent contacts.

To directly assess the contribution of αSyn’s C-terminal aromatic residues, we compared ^1^H–^15^N HSQC spectra of wild-type αSyn and a triple mutant (Y125G/Y133G/Y136G). The spectra were nearly identical (**Supplementary Fig. 11b**), indicating that the overall interaction topology of the optimized peptide remained unchanged despite the removal of these aromatic side chains. These results indicate that aromatic residues in both αSyn and the peptide promote condensate formation not only through π–π stacking but also via π–cation interactions involving nearby cationic residues. The combined contribution of these complementary forces is likely to enhance both the specificity and reversibility of condensate assembly.

Overall, peptide-induced αSyn phase separation is governed by a defined stoichiometry and enthalpy-driven interactions, with key contributions from aromatic, cationic, electrostatic, and hydrophobic forces that cooperatively stabilize the condensate network. Importantly, LLPS efficiency is not determined solely by the presence and affinity of detectable binding sites but also by cryptic or dynamic interactions that reinforce multivalent connectivity. The optimized peptide (FD1L_posi2_)_2_ exemplifies how rational sequence variation can strengthen such cooperative interactions, even when not all contact sites are directly observable by HSQC.

### Concentration-dependent interaction modes explain the bell-shaped phase diagram

To elucidate the molecular basis of the bell-shaped phase diagram, we examined the concentration dependence of peptide–protein interactions using NMR ^1^H–^15^N HSQC spectroscopy (**Fig. 5a** and **Supplementary Fig. 12**). We reasoned that within the condensed phase, αSyn and the peptide form a transient intermolecular network. Although these interactions are dynamic and fluid, they markedly restrict the rotational diffusion of αSyn, causing extensive line broadening across all residues such that NMR resonances become undetectable. Consequently, the observed signals primarily originate from αSyn in the dilute phase. Nevertheless, if αSyn exchanges between dilute and condensed phases on a μs–ms timescale, peak broadening may still reflect interactions occurring within the condensed phase. To exclude such contributions, we additionally recorded spectra of the supernatant following centrifugation (**Fig. 5b**).

**Fig. 5.**
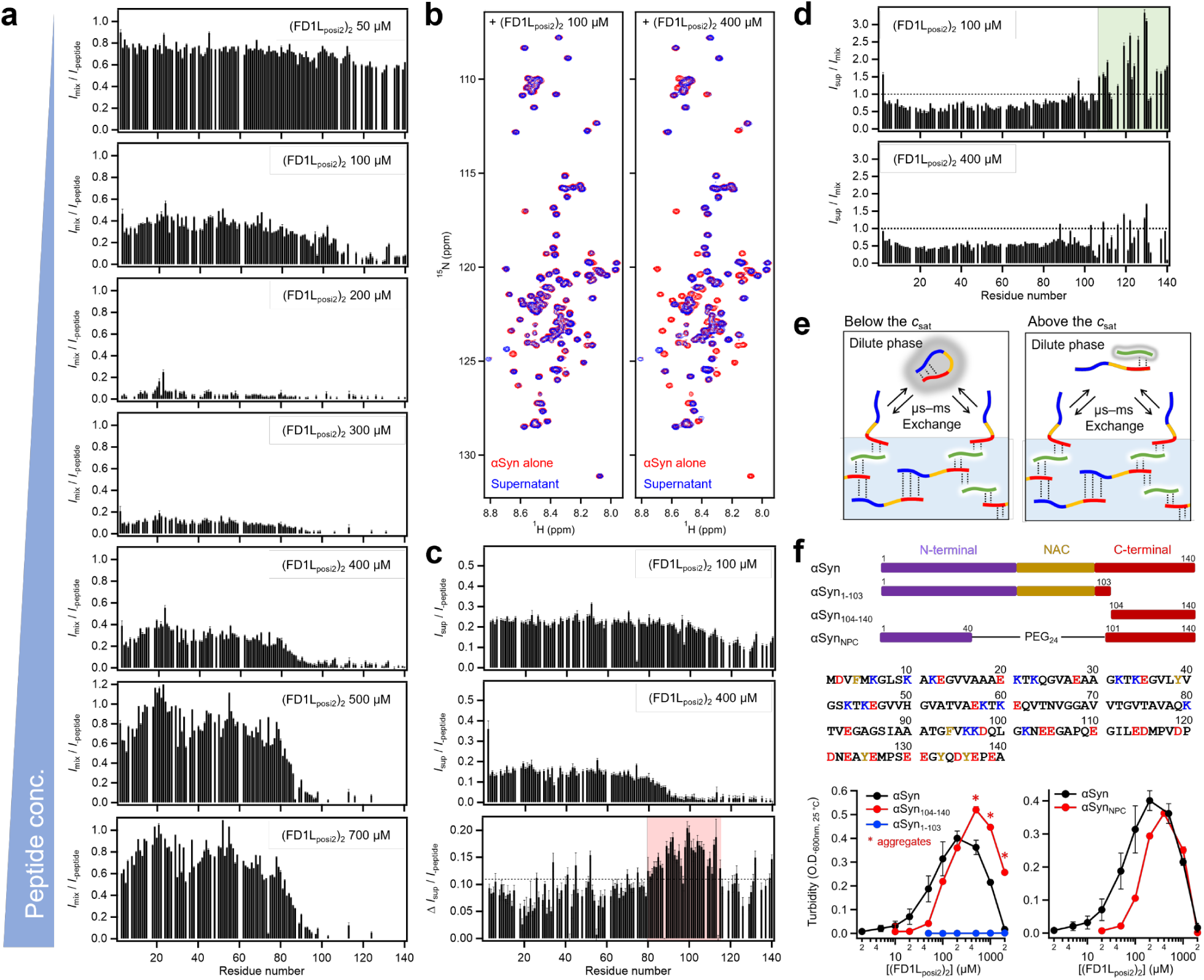
Interaction of the optimized peptides governing the bell-shaped phase diagram. **a** Changes in peak intensity ratios (*I*_+peptide_*/I*_-peptide_) from ^1^H-^15^N HSQC spectra of 100 μM αSyn at increasing peptide concentrations, showing progressive broadening and attenuation with increasing concentration. **b** Representative HSQC spectra at the low-concentration side of the bell-shaped diagram (100 μM peptide) and the high-concentration side (400 μM peptide), highlighting distinct interaction modes. **c** Residue-specific intensity ratios (*I*_sup_*/I*_-peptide_) comparing the low- and high-concentration regimes, together with difference maps (Δ *I*_sup_*/I*_-peptide_) illustrating the expanded interaction region at higher peptide concentrations. **d** Residue-specific intensity ratios (post-/pre-centrifugation) at 100 μM and 400 μM peptide concentrations, illustrating differences between droplet-internal dynamics and dilute-phase redistribution. **e** Schematic model of intermediate exchange of αSyn molecules at the droplet interface, with key interactions occurring at their C-terminal regions. **f** Schematic representation of αSyn domain-deletion constructs (C-terminal truncation αSyn_1–103_, N-terminal truncation αSyn_104–140_, and NAC-deletion mutant αSyn_NPC_) and their corresponding phase diagrams, demonstrating domain-specific contributions to LLPS efficiency and stability.

At sub-saturating peptide concentrations (e.g., 100 μM), we observed region-selective attenuation of peak intensities, predominantly in the C-terminal segment (residues 101–140). This broadening diminished in the supernatant spectrum (**Fig. 5a, c**, and green region in **Fig. 5d**), consistent with μs–ms exchange between dilute and condensed phases, where C-terminal contacts with the peptide in the condensed phase contribute to line broadening (**Fig. 5e**). Given that peptides themselves can dynamically shuttle between the two phases, transient encounters between αSyn and peptides in the dilute phase may also contribute to the observed signal attenuation. In either case, this region-specific effect supports the notion that the C-terminal domain drives partitioning into the condensed phase and directly mediates peptide-induced LLPS.

At the LLPS saturation point (∼200 μM), HSQC peak intensities dropped to ∼5% of their initial values. The residual signals closely matched the concentration of monomeric αSyn detected in the supernatant by HPLC (**Fig. 3i**), indicating that detectable signals could derive almost exclusively from a dilute phase of αSyn. This agreement confirms the sequestration of αSyn into a condensed phase, where it becomes NMR-invisible.

When the peptide concentration exceeded the saturation threshold (> 200 μM), marked broadening of C-terminal resonances was again observed in the supernatant. The similarity between pre- and post-centrifugation spectra (lower panel in **Fig. 5d**) indicates negligible contribution from condensed-phase species, suggesting the μs–ms dynamic interactions between peptide and αSyn within the dilute phase (**Fig. 5e**). Notably, the attenuation extended to residues 80–140 (red region in **Fig. 5c**), implying that an excess of peptide induces multivalent binding to both non-amyloid-β component (NAC) region, the aggregation-prone core of αSyn spanning residues 61–95, and C-terminal regions. These non-networking interactions inhibit productive intermolecular contacts, progressively dissolving condensates and dispersing peptide–αSyn complexes into the dilute phase. Since the peak intensities from the N-terminal region nearly returned to the levels observed before peptide addition, most αSyn molecules are inferred to have diffused from the condensed phase into the dilute phase.

To assess the functional relevance of individual regions, we examined truncated αSyn variants (**Fig. 5f**). The C-terminal deletion mutant (αSyn_1–103_) failed to undergo LLPS, confirming the essential role of the C-terminal region. The N-terminal deletion construct (αSyn_104–140_) retained LLPS capability but exhibited peptide-induced aggregation above 500 μM. By contrast, the NAC-deletion variant (αSyn_NPC_, residues 41–100 replaced with a flexible linker) showed moderately reduced LLPS efficiency yet maintained a bell-shaped phase diagram. Together, these results establish the C-terminal region as the primary determinant of LLPS, with the NAC region modulating phase separation efficiency and the N-terminal region influencing droplet fluidity.

The above results indicate that the bell-shaped phase behavior arises from a concentration-dependent switch in interaction modes. Below the critical peptide concentration, condensate formation is driven predominantly by C-terminal interactions. Beyond of this threshold, additional contacts—particularly around residues 101–110—disrupt intermolecular network connectivity, promoting dispersion of peptide–αSyn complexes into the dilute phase. This mechanistic transition explains both the emergence and collapse of LLPS across the peptide concentration range.

### LLPS-inducing peptides suppress αSyn fibril elongation through C-terminal binding

To investigate how LLPS-inducing peptides influence amyloid fibril formation of αSyn, we first examined spontaneous fibrillation under shaking conditions in the absence of preformed seeds (**Fig. 6a**). In these assays, the presence of low concentrations of the optimized peptide (FD1L_posi2_)_2_ (1–20 μM) markedly accelerated the increase in Thioflavin T (ThT) fluorescence intensity, indicating enhanced nucleation of αSyn fibrils. This acceleration likely arose from two cooperative effects: local concentration of αSyn molecules within transient peptide-induced condensates, and partial exposure of the NAC region within these dense phases, which facilitates intermolecular contacts required for nucleation of amyloid fibril formation^36,43–45^.

**Fig. 6.**
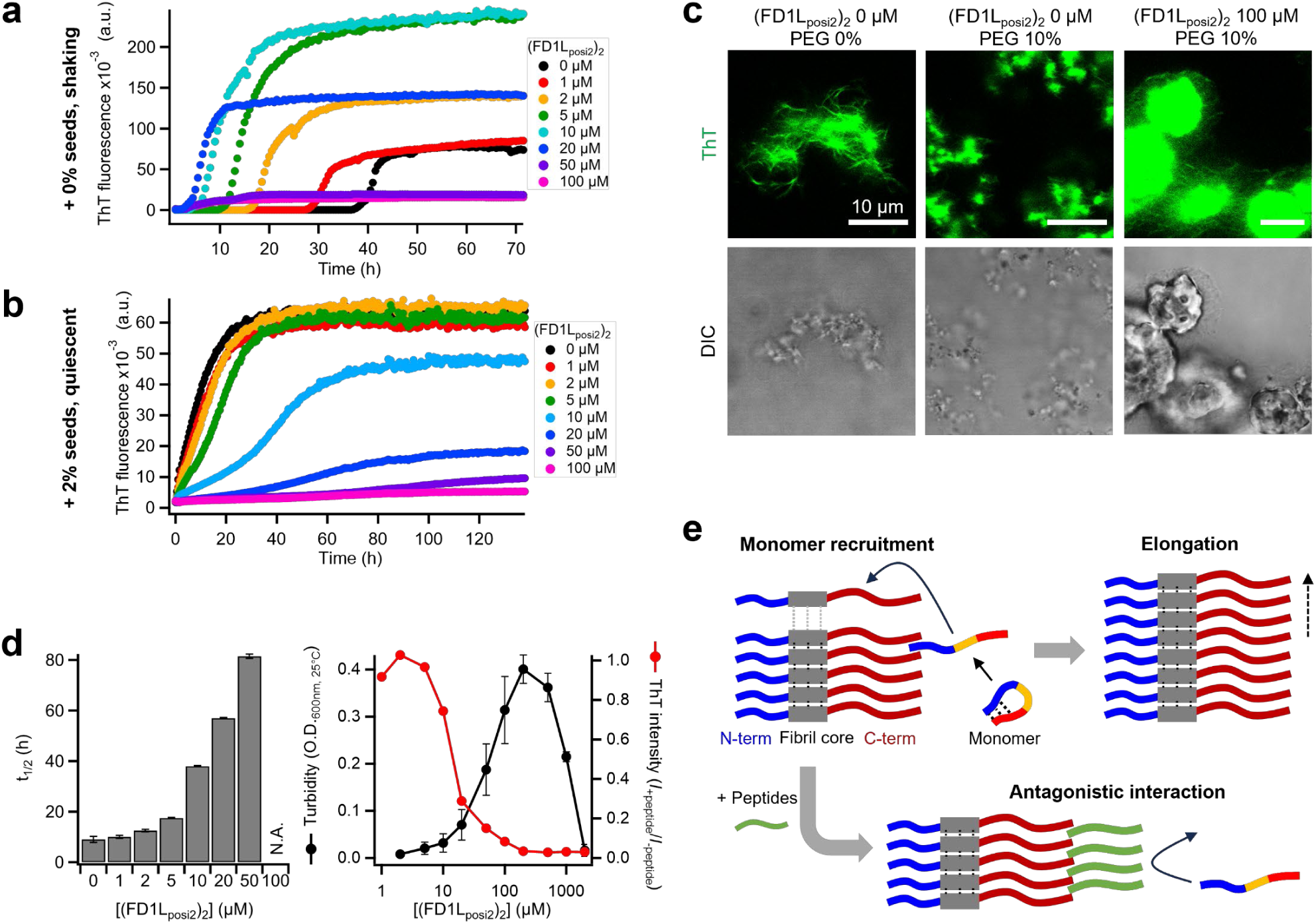
Biphasic effects of LLPS-inducing peptides on αSyn fibril formation. **a** Spontaneous fibrillation of αSyn monitored by ThT fluorescence under shaking conditions. Low concentrations of the optimized peptide (FD1L_posi2_)_2_ (1–20 μM) markedly accelerated the nucleation of αSyn fibrils. **b** Seeded fibril elongation assays performed in the presence of 2 % preformed αSyn fibrils. Addition of (FD1L_posi2_)_2_ suppressed ThT fluorescence intensity and slowed elongation in a concentration-dependent manner. **c** Representative fluorescence (ThT) and DIC microscopy images obtained after completion of the seeding experiments, illustrating the aggregates at 100 μM (FD1L_posi2_)_2_. **d** Comparison of the lag time of the αSyn fibril formation (*t*_1/2_, defined as the time to reach half-maximal ThT intensity) and endpoint ThT fluorescence intensity with the peptide-induced LLPS phase diagram. Fibril inhibition is observed even under conditions where droplets are not formed, suggesting an LLPS-independent mechanism of inhibition. **e** Schematic model illustrating the proposed mechanism by which (FD1L_posi2_)_2_ inhibits fibril elongation. The peptide binds to the C-terminal region of αSyn monomers, preventing their recruitment to the surface of growing fibrils and thereby blocking elongation.

We next evaluated the effect of LLPS-inducing peptides on the amyloid fibril formation of αSyn by performing ThT fluorescence assays using 2% preformed αSyn fibrils as seeds (**Fig. 6b, c, Supplementary Fig. 13**). In the presence of (FD1L_posi2_)_2_, both ThT fluorescence intensity and fibril elongation rate was suppressed in a concentration-dependent manner (**Fig. 6d**). Notably, endpoint ThT intensities at each peptide concentration correlated with the LLPS phase diagram, yet fibril inhibition was also observed under peptide-rich conditions where droplet formation did not occur. These results indicate that suppression of fibril formation is not solely attributable to LLPS but likely arises from direct interactions of the peptide with the C-terminal region of αSyn.

Previous studies have established that αSyn monomers are recruited to the surface of growing fibrils through C-terminal interactions^46^—the same region targeted by (FD1L_posi2_)_2_. Thus, it is plausible that the peptide competitively interferes with monomer binding, thereby blocking recruitment and subsequent fibril elongation (**Fig. 6e**). At lower peptide concentrations (< 20 μM), where LLPS occurs but is not saturated, modest changes in ThT fluorescence were observed. Because ThT fluorescence is sensitive not only to fibril quantity but also to morphology and packing, these changes may reflect alterations in fibril structure or supramolecular organization^47,48^ rather than complete inhibition of elongation^49^.

Taken together, these results reveal a biphasic influence of the peptide on αSyn fibrillation. At low concentrations, peptide-induced LLPS accelerates nucleation through condensate-driven molecular concentration and exposure of the NAC region, whereas at high concentrations, excess peptide suppresses elongation by competitively binding to αSyn and preventing productive assembly. Thus, (FD1L_posi2_)_2_ exemplifies how LLPS-modulating peptides can switch between promoting and inhibiting fibril formation depending on concentration, offering mechanistic insight into the dual relationship between phase separation and amyloidogenesis.

## Discussion

Understanding and engineering the rules that govern liquid–liquid phase separation (LLPS) of intrinsically disordered proteins is essential for controlling biomolecular condensates in vitro and in vivo. In this study, we established a systematic framework for designing and optimizing de novo peptides that induce and modulate the LLPS of α-synuclein (αSyn). Building upon our previous identification of FL2 and FD1, we demonstrated that LLPS efficiency can be dramatically enhanced through the rational sequence optimization guided by deep mutational scanning (DMS) and biophysical characterization. This work highlights how peptide sequence features—including binding affinity, solubility, and multivalency—synergistically determine condensate formation and material properties.

Our results reveal several key principles for the molecular control of LLPS. First, optimization of peptide sequences under physiologically relevant conditions showed that affinity-enhancing substitutions and solubility tuning are critical for balancing droplet formation with the avoidance of nonspecific aggregation. The identification of optimized peptides, particularly FD1L_posi2_, demonstrated that high LLPS-inducing efficiency can be achieved without compromising solubility or target specificity. Moreover, tandem-repeat peptides further amplified LLPS activity through multivalency, although often at the cost of reduced droplet fluidity. Notably, FD1L_posi2_ uniquely retained both high efficiency and fluidity, enabling construction of a concentration-dependent phase diagram. Importantly, the application of DMS not only clarified sequence–activity relationships but also provided a powerful framework for guiding rational peptide design, suggesting that similar approaches could broadly accelerate the development of optimized LLPS modulators with tailored physicochemical properties.

Mechanistic analyses revealed that peptide–protein interactions are concentration dependent and underlie the characteristic bell-shaped phase diagram depicted in **Figure 7**. At sub-saturating concentrations, peptides were largely sequestered into the condensed phase, leaving ∼2 μM peptide in the dilute phase, while αSyn interacted with them in a 2:1 stoichiometry (αSyn: (FD1L_posi2_)_2_). This multivalent binding promoted networked interactions that drive condensate formation. At the saturation point, residual αSyn remained at ∼5 μM in the dilute phase. Above this critical concentration, however, excess peptides established non-networking contacts that disrupted condensate connectivity. Detailed thermodynamic and structural analyses further established that LLPS arises from a cooperative balance among electrostatic, aromatic, π–cation, and hydrophobic interactions. Salt-dependent suppression of phase separation revealed that electrostatic attraction between the highly anionic C-terminal region of αSyn and cationic residues within the peptide is indispensable for condensate stability. Conversely, 1,6-hexanediol, which weakens hydrophobic interactions, also reduced LLPS efficiency, underscoring the contribution of hydrophobic contacts to network cohesion. Mutational and NMR analyses demonstrated that π–π interactions between peptide residues Y1, W4, W8, and F9 and αSyn residues Y125, Y133, and Y136 are central to condensate stabilization. These aromatic contacts, together with π–cation interactions involving neighboring cationic residues, likely underlie the high target specificity and reversible multivalency characteristic of FD1-derived peptides. Enhanced LLPS efficiency by optimized peptides further stems from long-range electrostatic complementarity that reinforces the condensate network.

**Fig. 7.**
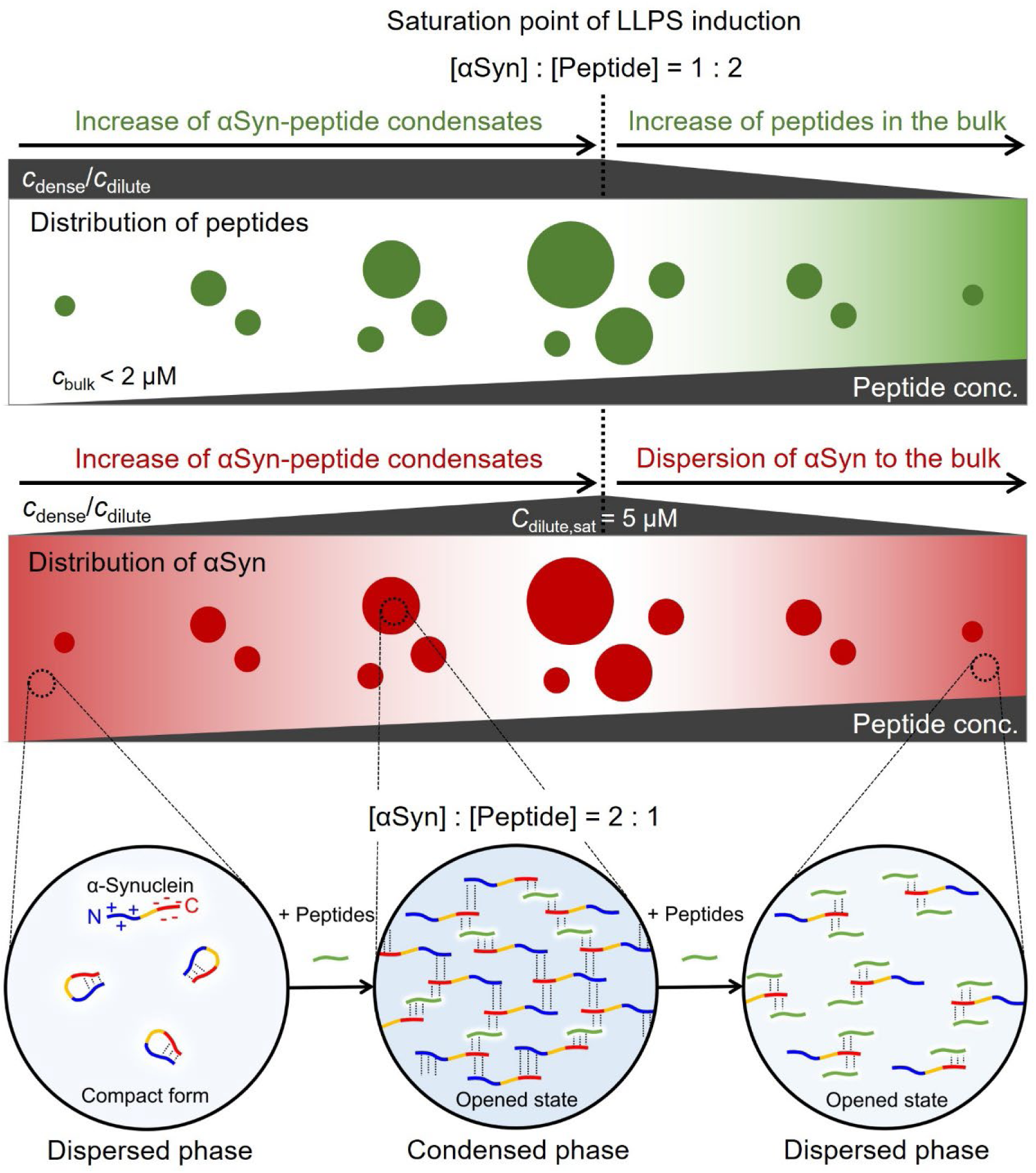
Proposed mechanistic model explaining the bell-shaped phase diagram of αSyn condensates modulated by optimized LLPS-inducing peptides. Under physiological conditions, αSyn adopts a compact conformation stabilized by long-range intramolecular interaction between the N- and C-terminal regions^44,67–69^. At concentrations below the LLPS critical point, (FD1L_posi2_)_2_ binds selectively to the C-terminal region of αSyn in a [αSyn] : [peptide] = 2 : 1 stoichiometry, promoting multivalent intermolecular interactions that drive phase separation and droplet formation^36,55^. Nearly all peptide molecules are incorporated into the condensed phase, while a small residual fraction (∼2 μM) remains in the dilute phase. As peptide concentration increases and approaches the critical threshold ([αSyn] : [peptide] = 1 : 2), droplet formation saturates and the network of interactions reaches its maximum, with αSyn remaining at ∼5 μM in the dilute phase. Beyond this saturation point, excess peptide molecules engage in additional non-productive interactions with αSyn monomers, likely occluding binding interfaces and preventing intermolecular bridging. This shifts the interaction network balance, leading to dispersal of both peptide and αSyn into the dilute phase and dissolution of condensates.

At high peptide concentrations, new interactions emerged within αSyn residues 101–110 (including K102, E104, E105, and E110), forming non-productive complexes that redistributed both peptide and protein into the dilute phase. This mechanistic shift is well consistent with the explanation where the LLPS efficiency on peptide concentration showed bell-shaped dependent profile, illustrating how a balance of attractive and competing interactions governs condensate stability.

LLPS-mediated liquid-to-solid phase transitions into pathological aggregates have been reported for various intrinsically disordered proteins (IDPs)^50,51^. Consistent with these observations, we found that LLPS-inducing peptides exert a biphasic influence on αSyn fibrillation. At low peptide concentrations (1–20 μM), (FD1L_posi2_)_2_ accelerated spontaneous fibril nucleation, likely due to local enrichment of αSyn^8,52–54^ and exposure of its aggregation-prone NAC region^36,45,55^ within transient condensates. In contrast, at higher concentrations, the same peptide suppressed seeded fibril elongation in a concentration-dependent manner by competitively binding to the C-terminal region of αSyn and preventing monomer recruitment to fibril ends. Thus, the peptide promotes nucleation at low concentrations but inhibits elongation at high concentrations, illustrating how LLPS-modulating peptides can reversibly shift the balance between condensation and aggregation.

Beyond αSyn, our results also highlight the utility of DMS as a general strategy for peptide optimization. By quantitatively mapping the contribution of each residue to binding and condensate formation, DMS provides a powerful framework not only for discovering LLPS-inducing sequences but also for fine-tuning pre-designed, target-specific peptides^56,57^. This approach enables systematic improvement of affinity, solubility, and specificity, and is likely to accelerate the development of tailored LLPS modulators across diverse protein systems.

In conclusion, our study provides mechanistic insights and generalizable design principles for peptide-based modulation of LLPS. The interplay of π–π contacts, long-range electrostatics, solubility, multivalency, and concentration-dependent interaction modes forms the foundation for creating next-generation LLPS modulators with defined physicochemical properties and molecular specificity. These findings offer a roadmap for tailoring condensate behavior in diverse protein systems, with potential applications ranging from the mechanistic dissection of cellular condensates to the development of therapeutics targeting pathological phase transitions.

## Materials and Methods

### Recombinant expression and purification of αSyn

The plasmid that expresses human αSyn and the mutants were amplified as previously described^58^. αSyn, αSyn_1-103_, and αSyn_104-140_ were expressed in an *Escherichia coli* BL21(DE3) transformed by pET-αSyn in 2 L flasks at 37 °C with 1 L of Luria-Bertani (LB) medium. Isotopically labeled ^15^N-αSyn were expressed in 2 L flasks at 37 °C with 1 L of minimal M9 batch medium. Cells were suspended in purification buffer (50 mM Tris-HCl, pH 7.5, containing 1 mM EDTA and 0.1 mM dithiothreitol), disrupted using sonication, and centrifuged (10,000×*g*, 30 min). Streptomycin sulfate (final 5%) was added to the supernatant to remove nucleic acids. After removal of nucleic acids by centrifugation, the supernatant was heated to 80 °C for 30 min and then centrifuged. In this step αSyn remained in the supernatant. The supernatant was precipitated by the addition of solid ammonium sulfate to 70% saturation, centrifuged, and dialyzed overnight and then applied onto a HiTrap-Q column (cytiva) with 50 mM Tris-HCl buffer, pH 7.5, containing 1 mM EDTA and 0.1 mM dithiothreitol as running buffer. Samples were eluted with a linear gradient of 0–1 M NaCl. Collected fractions were dialyzed overnight and then applied onto a reversed-phase HPLC (RP-HPLC), using a Prominence HPLC system (Shimadzu) under linear gradient conditions. Mobile phase A (comprising water with 0.1% TFA) was mixed with mobile phase B (0.1% TFA in acetonitrile). Purified peptides were lyophilized, and molecular mass was confirmed by matrix-assisted laser desorption/ionization time-of-flight mass spectroscopy (MALDI MS), using an UltrafleXtreme instrument (Bruker Daltonics).

### αSyn fibrils for deep mutational scanning

Since polymorphism is a characteristic property of amyloid fibrils^59^, it is important to prepare and apply homogeneous fibril for the deep mutational scanning. We first amplified specific αSyn fibril by repeating seeding experiment. Solutions of monomeric αSyn were prepared by dissolving the lyophilized αSyn with PBS buffer. Solutions were filtered using a 0.22 μm PVDF filter, and the αSyn concentration was determined by NanoDrop using ε_280_ = 5120 L mol^−1^ cm^−1^. Seeding experiments were performed by adding 5% (v/v) preformed fibrils to 100 μM monomeric αSyn solution. First generation of fibrils for seeding experiments were prepared by spontaneous fibril formation by monitoring ThT fluorescence. Assays were initiated by placing the 96-well plate at 37 °C with a cycle of 3 min shaking and 27 min quiescence in a plate reader (Flex station; Molecular Devices). Preformed fibrils were well fragmented by ultrasonication before seeding, and the seeded solution was incubated at 37 °C for 1 week. This seeding experiment was repeated six times with PBS buffer at pH 7.5. The homogeneity of the morphology of the 7th generation of amyloid fibrils was confirmed by analyzing 10 AFM images. We thus decided to use this particular sample for our RaPID campaign._In order to immobilize the fibrils on Dynabeads, human (His)_6_-αSyn (Wako Pure Chemical Industries Ltd) was attached to fibril ends in an amyloid propagation manner by adding 5 % of (His)_6_-αSyn to the 7th generation of fibrils.

### Deep mutational scanning of peptides

Deep mutational scanning of peptides targeting human αSyn fibrils were performed using the RaPID as previously reported^49^ slightly modified as follows: mRNA library containing a single ‘NNK’ degenerate codon at different positions in the FL2 and FD1 encoding region was used. To generate the peptide library initiated with Tyr, mRNA–puromycin library was translated in a methionine-deficient FIT system^60^ with Tyr-tRNA^fMet^_CAU_. The buffer-exchanged peptide-mRNA fusions were collected and 5 μL of blocking buffer (PBST containing 0.2% acetylated BSA; Life Technologies) was added. The peptide-oligonucleotides (mRNA/cDNA) fusions were then incubated with Dynabeads His-tag isolation and pulldown (invitrogen) for 30 min at room temperature (negative selection against beads). A 2.5 μL aliquot of the peptide-mRNA fusions was taken from the mixture and saved for the determination of the total amount of input DNA. The unbound fraction was then incubated with human αSyn fibrils immobilized on Dynabeads for 30 min. αSyn fibrils were masked with 2 mg/mL yeast tRNA (invitrogen) in advance of applying the library. During selection, αSyn fibrils were treated at room temperature to avoid cold denaturation^42^. The resultant complementary DNAs (output DNA) were eluted by mixing with 1 × PCR reaction buffer and heating at 95 °C for 5 min, followed by immediate separation of the supernatant from the beads. A small fraction of the DNA (input DNA and output cDNA) was allocated to real-time PCR quantification using a LightCycler 2.0 (Roche); the remainder was amplified by PCR. After precipitation, the observed enrichments were subjected to further DNA deep sequencing using the MiSeq sequencing system (Illumina). All DNA oligos were purchased from eurofins Genomics and are listed in Table S1. Equations [1-3] were used to calculate the enrichment score *E* for each peptide. An *E* = 1 means identical enrichment to the wild-type peptide.

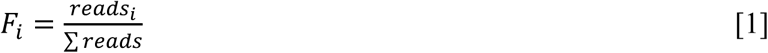

*F_i_*: Fraction of reads for peptide *i* (wild-type or mutant) relative to the total number of reads (wild-type and all mutants).

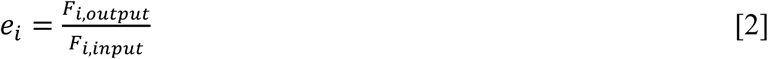

*e_i_*: Raw enrichment score for mutant *i*, calculated as the ratio of its fractional abundance in the output library (*F_i,output_*) to that in the input library (*F_i,input_*).

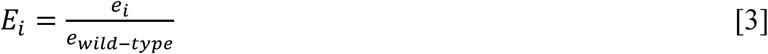

*E_i_*: Normalized enrichment score for mutant *i*, where values greater or less than 1 indicate higher or lower enrichment relative to the wild type, respectively.

### Chemical synthesis of peptides

Macrocyclic and liner peptides were synthesized by standard Fmoc solid-phase peptide synthesis (SPPS) using a Syro Wave automated peptide synthesizer (Biotage). The resulting peptide–resin (25 μmol scale) was treated with a solution of 92.5% trifluoroacetic acid (TFA), 2.5% water, 2.5% triisopropylsilane and 2.5% ethanedithiol, to yield the free linear *N*-ClAc-peptide. Following diethyl ether precipitation, the pellet was dissolved in 10 ml triethylamine containing DMSO and incubated for 1 h at 25 °C, to yield the corresponding macrocycle. The peptide suspensions were then acidified by addition of TFA to quench the macrocyclization reaction. The macrocycle was purified by RP-HPLC, using a Prominence HPLC system (Shimadzu) under linear gradient conditions. Mobile phase A (comprising water with 0.1% TFA) was mixed with mobile phase B (0.1% TFA in acetonitrile). Purified peptides were lyophilized *in vacuo* and molecular mass was confirmed by MALDI MS, using an Autoflex II instrument (Bruker Daltonics). Theoretical scores of water solubility of linear peptides were determined using the CamSol method^39^. Tandem-repeat peptides were synthesized only up to two repeats due to the limitations of solid-phase peptide synthesis.

For the fluorescein- and Cy5-labelling of peptides, peptide-Cys and peptide-Lys was synthesized by Fmoc SPPS. The obtained peptides were treated with the six equivalents of 5-(Iodoacetamido)fluorescein (Santa Cruz Biotechnology), or Cy5-NHS (BroadPharm) in DMSO. The resulting peptides were purified by RP-HPLC and lyophilized *in vacuo*.

### Phase separation assay

Monomeric αSyn solutions were prepared by dissolving lyophilized αSyn in 10 mM NaOH and adjusting to neutral pH. The solutions were filtered through a 0.22 μm filter, and αSyn concentrations were determined using a NanoDrop spectrophotometer. The monomer was diluted with water to 400 μM and stored at −80 °C. Phase separation was induced by mixing αSyn (in the desired buffer, pH 7.5) with 10% PEG and peptides dissolved in DMSO. The peptide stock was diluted to achieve a final DMSO concentration of 1%. Differential interference contrast (DIC) images were acquired at room temperature using a Leica DMI6000 B microscope with a 40× objective. Images were recorded at 696 × 520 pixels with 24-bit depth. Unless otherwise indicated, the αSyn concentration was fixed at 100 μM.

For turbidimetry, 100 μM αSyn in the presence of 10% PEG was incubated with various peptide concentrations for 15 min at 4 °C. Measurements were performed using a Jasco V-670 spectrometer (JASCO) at 600 nm. Temperature was controlled with a Peltier unit (JASCO) and a 1 mm path-length cuvette. Quantification of αSyn and peptide in the dilute phase was performed by UPLC (Shimadzu). After centrifugation (50 μL, 11,000 × g, 10 min), 20 μL of the supernatant was injected into a reversed-phase column.

For confocal microscopy, mixtures of 1% (FD1L_posi2_)_2_-Fluor or (FD1L_posi2_)_2_-Cy5 with 99% unlabeled (FD1L_posi2_)_2_ were used as LLPS-inducing peptides for αSyn. Wild-type αSyn was mixed with αSyn-Rhod at a 99:1 molar ratio. To minimize the effects of (FD1L_posi2_)_2_-Fluor and (FD1L_posi2_)_2_-Cy5 on droplet liquidity, their concentrations were reduced by mixing with unlabeled (FD1L_posi2_)_2_.

### Confocal microscopy

The fusion event of αSyn liquid droplets in vitro was visualized with a using a Leica TCS SP8 confocal microscope with a 63× oil objective lens at room temperature. Rhodamine-labeled αSyn, fluorescein- or Cy5-labeled peptide, and Thiolfavin T (ThT) (Wako Pure Chemical Industries Ltd.) were observed using appropriate fluorescence channels (488 nm for fluorescein, 561 nm for rhodamine and Cy5, and 442 nm for ThT). All the images were captured at a resolution of 512×512 pixels at 24-bit depth. The FRAP measurements were performed using a Leica TCS SP8 confocal microscope. A region of interest (ROI) with a radius of 1.0 μm was bleached using an appropriate laser, and the recovery of the bleached spots was recorded using the software provided with the instrument. The fluorescence recovery was background-corrected, normalized, and plotted using Igor Pro.

### Isothermal titration calorimetry (ITC)

ITC measurements were performed to study the binding between αSyn synthesized peptides using a Nano ITC instrument (TA Instruments). The peptides were dissolved in DMSO and a 70 mM (FD1L_posi2_)_2_ stock solution was prepared. A 200 μM (FD1L_posi2_)_2_ in PBS buffer (1% DMSO, 10% PEG) was then injected into the sample cell containing approximately 190 μl of αSyn monomer solution at 50 μM in PBS (1% DMSO, 10% PEG). ITC titrations were carried out at 25 °C with 2.5 μl injections for a total of 18 injections with stirring at 400 rpm. The data were fitted using an independent one-binding site model.

### NMR measurements

All NMR experiments were performed on Bruker BioSpin Avance III HD spectrometers with TCI triple-resonance cryogenic probe-head with basic ^1^H resonance frequencies of 599.96 and 800.23 MHz. ^1^H-^15^N HSQC spectra of 0.1 mM [^15^N]-αSyn titrated with LLPS-induce peptides at different concentrations were measured in PBS buffer (pH 7.4), 10% (v/v) PEG8000, 5% (v/v) D_2_O, and 1% (v/v) DMSO at 4 °C. Three-dimensional (3D) spectra for main-chain signal assignments: HNCACB, CBCA(CO)NH, HNCA, HN(CO)CA, HNCO, HN(CA)CO and HN(CA)NNH were acquired for 0.3 mM [^13^C, ^15^N]-αSyn in the same buffer condition as that in the peptide-titration experiments. All 3D spectra were acquired by means of non-uniform sampling (NUS) to randomly reduce t1 and t2 time-domain data points around to 25%. The uniformly sampled data were reconstructed from the raw NMR data according to the sparse sampling schedules using techniques such as IST and SMILE^61–63^. All NMR package of NMR tools named MagRO-NMRViewJ, version 2.01.39 [the upgraded version of Kujira^64^], on NMRView^65^ was used for the signal assignment, and Sparky for the signal intensity estimation ^66^.

### Fluorescence assay

The αSyn monomer was diluted to the desired concentration with a 10×PBS buffer and supplemented with 20 μM ThT from a 1 mM stock. All samples were prepared in low-binding Eppendorf tubes on ice. Each sample was then pipetted into multiple wells of a 96-well half-area, low-binding polyethylene glycol coating plate (Corning 3881) with a clear bottom, at 80 μl per well. Assays were initiated by placing the 96-well plate at 37 °C with a cycle of 3 min shaking and 27 min quiescence in a plate reader (Flex station; Molecular Devices). The fluorescence of ThT was measured through the bottom of the plate with a 440 nm excitation filter and a 480 nm emission filter, with three repeats per sample.

## Acknowledgements

This work was supported by JST, ACT-X Grant Number JPMJAX2113, Japan to T.I., the Japan Society for the Promotion of Science (JSPS) Grant-in-Aid for Transformative Research Areas (JP25H02246) to T.I., Astellas Foundation for Research on Metabolic Disorders to T.I., The Sumitomo Foundation to T.I., the Technology Licensing Fund generated by H.S., and JSPS Grant-in-Aid for Specially Promoted Research (JP20H05618) to H.S..

## Author Contributions

T. Ikenoue and H.S. were involved in the design of research. T. Ikenoue performed all in vitro experiments. T.K. and T. Ikegami contributed to NMR measurements. T. Ikenoue and H.S. wrote the paper. All authors discussed the results and commented on the manuscript.

