## Supplementary information for "Fine-Tuning α-Synuclein Phase Separation through Sequence-Optimized Peptide Modulators"

**Supplementary Table 1. Oligonucleotides**

| Name | Sequence |
|---|---|
| FL2L wild-type DNA | CTTTAAGAAGGAGATATACATATGCAGAGGTGGGGTAAGCCTCGGTGG TGGCGTACTTCTTATTGGTCTGGCAGCGGCAGCGGCAGC |
| FD1L wild-type DNA | CTTTAAGAAGGAGATATACATATGCTTGGTTGGCATAAGTTGTGGTTTC AGCCGTATGGTCGTTCTGGTGGCAGCGGCAGCGGCAGC |
| Puromycin construct | pCTCCCGCCCCCCGTCC–PEG linker–CC–puromycin |
| PCR primers used for deep mutational scanning | TTTCCGCCCCCCGTCCTAGCTGCCGCTGCCGCTGCCGCA |
| | TAATACGACTCACTATAGGGTTAACTTTAAGAAGGAGATATACATA |

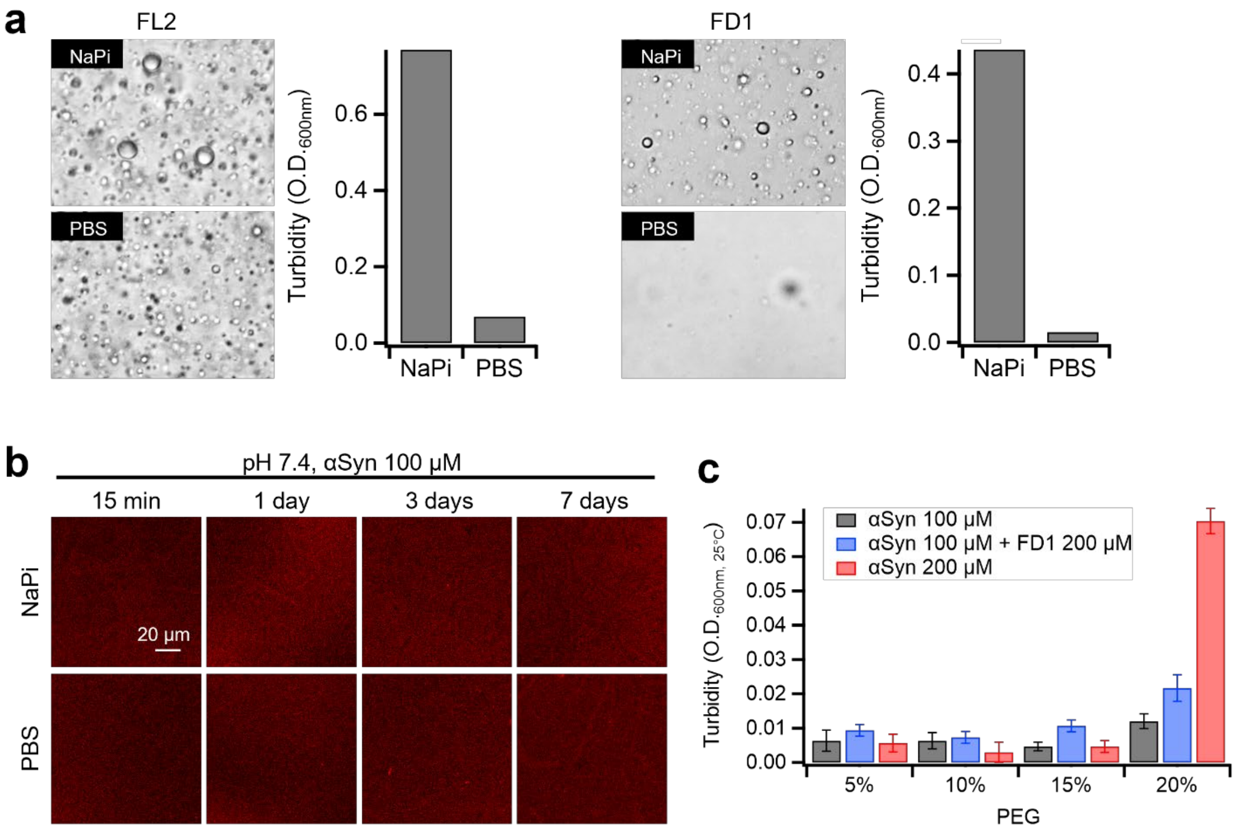


**Supplementary Fig. 1: Elevated salt concentrations reduce droplet formation induced by LLPS-promoting peptides.**

**a** Comparison of the LLPS efficiency in 20 mM sodium phosphate (NaPi) buffer versus PBS (10 mM NaPi, 140 mM NaCl) in the presence of LLPS-inducing peptide FD1 and FL2. **b** In the absence of peptides, αSyn does not form condensate even after one week of incubation. **c** Increasing concentration of PEG promote LLPS within 30 min.

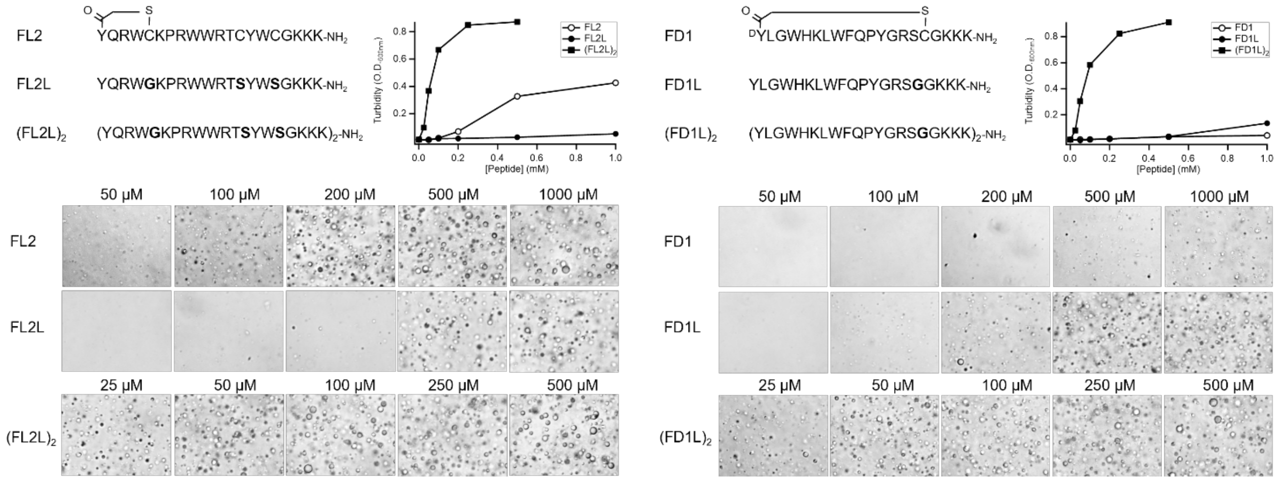


**Supplementary Fig. 2: Structural conversion of peptides from cyclic to linear forms alters LLPS efficiency.**

To eliminate the influence of cysteine side chains used for cyclization, cysteine residues were substituted with glycine. Additional cysteines were replaced with serine to prevent peptide multimerization. Sequence-repeated linear peptides, $(FL2L)_2$ and $(FD1L)_2$, exhibited markedly enhanced LLPS efficiency.

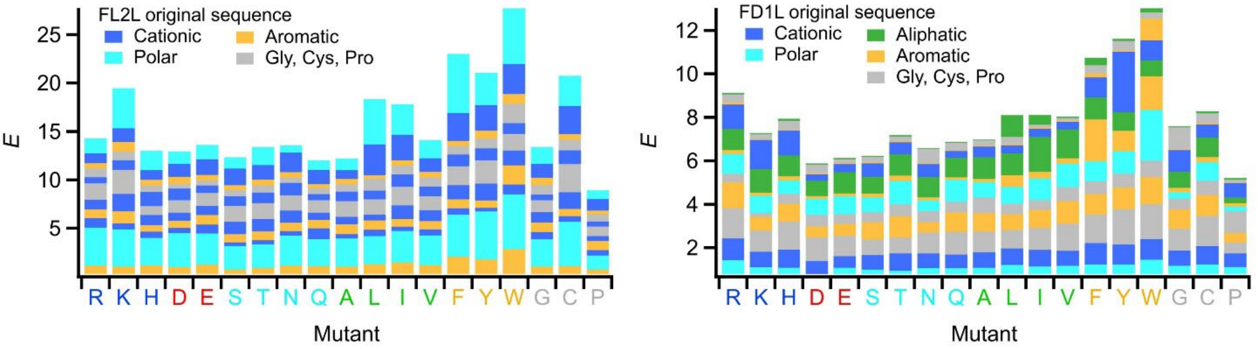


**Supplementary Fig. 3: Global trends in affinity gains from deep mutational scanning of LLPS-inducing peptides.**

Bar plots summarize the cumulative enrichment scores of single-point mutants derived from FL2L and FD1L scaffolds. The x-axis represents the amino acid residue introduced by mutation, and the y-axis shows the summed enrichment score, reflecting the overall affinity gain relative to the parental sequence. This analysis enables comparison of how substitution with different amino acids influences αSyn binding affinity. Mutants are color-coded according to the chemical class of the original residue in the parent peptide: cationic, aliphatic, polar, aromatic, or other.

| Peptide | Sequence | Solubility score | Peptide | Sequence | Solubility score |
|---|---|---|---|---|---|
| FL2L | YQRWGKPRWWRTSYWSGKKK-$NH_2$ | 1.998 | FD1L | YLGWHKLWFQPYGRSGGKKK-$NH_2$ | 1.377 |
| FL2LΔ | YQRWGKPRWWRTSYWS-$NH_2$ | 1.435 | FD1LΔ | YLGWHKLWFQPYGRSG-$NH_2$ | 0.742 |
| FL2L$_{arom}$ | YFRWGKWRWWRYSYWSGKKK-$NH_2$ | 1.167 | FD1L$_{arom}$ | YLGWYKLWFWPYYRSGGKKK-$NH_2$ | 0.197 |
| FL2L$_{posi}$ | YKRWGKKRWWRRSYWSGKKK-$NH_2$ | 2.420 | FD1L$_{posi}$ | YLGWHKKWFQPRGRRGGKKK-$NH_2$ | 2.444 |
| FL2L$_{pol}$ | YTRWGKSRWWRNQYWSGKKK-$NH_2$ | 2.018 | FD1L$_{pol}$ | YLNWHKTWFQPYQRSGGKKK-$NH_2$ | 1.586 |
| FL2$_{arom2}$ | YFRWWKPRWWRYSYWSGKKK-$NH_2$ | 1.062 | FD1L$_{posi2}$ | YLGWHKLWFQPYRRRGGKKK-$NH_2$ | 1.796 |
| FL2L$_{posi2}$ | YKRWGKPRWWRRSYWSGKKK-$NH_2$ | 2.300 | FD1L$_{pol2}$ | YLNWHKLWFQPYQRSGGKKK-$NH_2$ | 1.415 |
| FL2L$_{pol2}$ | YTRWGKPRWWRNSYWSGKKK-$NH_2$ | 2.057 | JMD$_{83-96}$ | KLKRKYWWKNLKMM-$NH_2$ | 2.142 |

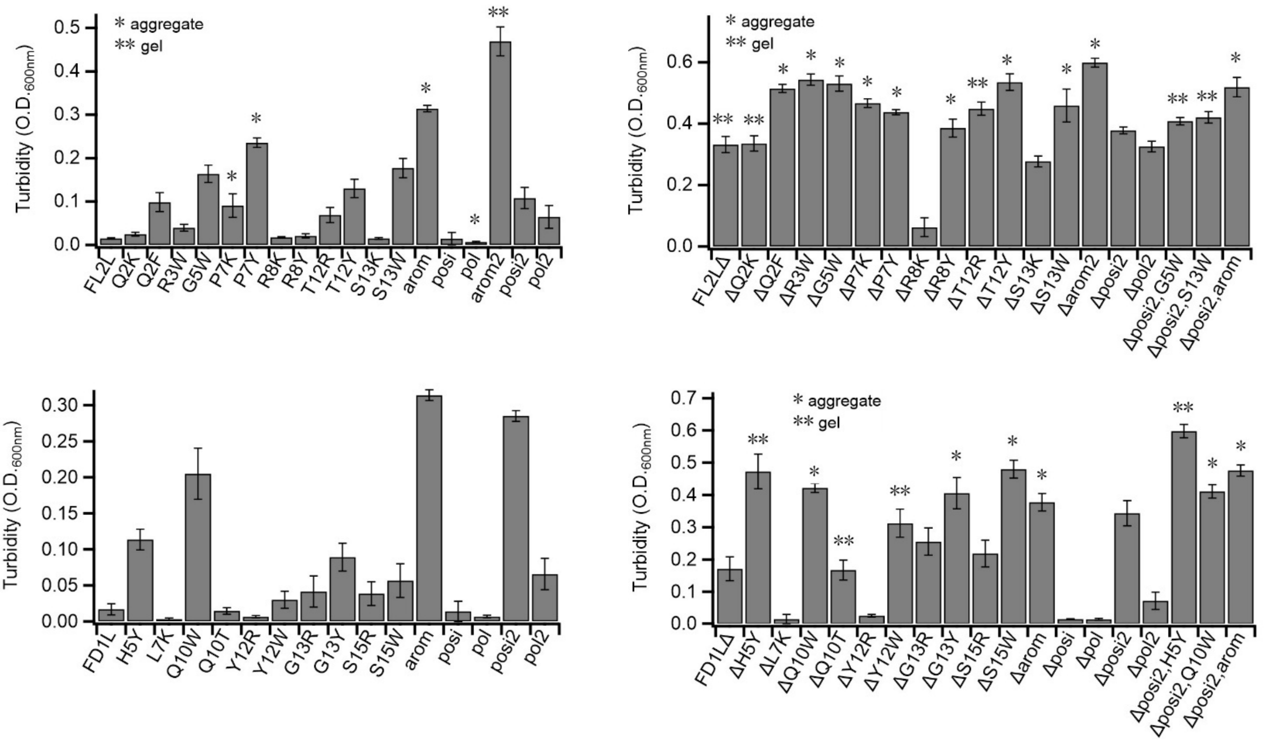


**Supplementary Fig. 4: Amino acid sequences of multiply mutated peptides and the impact of mutations on LLPS efficiency.**

Solubility scores were estimated using the CamSol method. For comparison, the sequence of JMD83–96 is also shown. LLPS efficiency of each selected variant was assessed by turbidimetry. The physical state of the resulting assemblies—liquid-like droplets, gel-like structures, or aggregates—was determined based on morphology observed in DIC imaging.

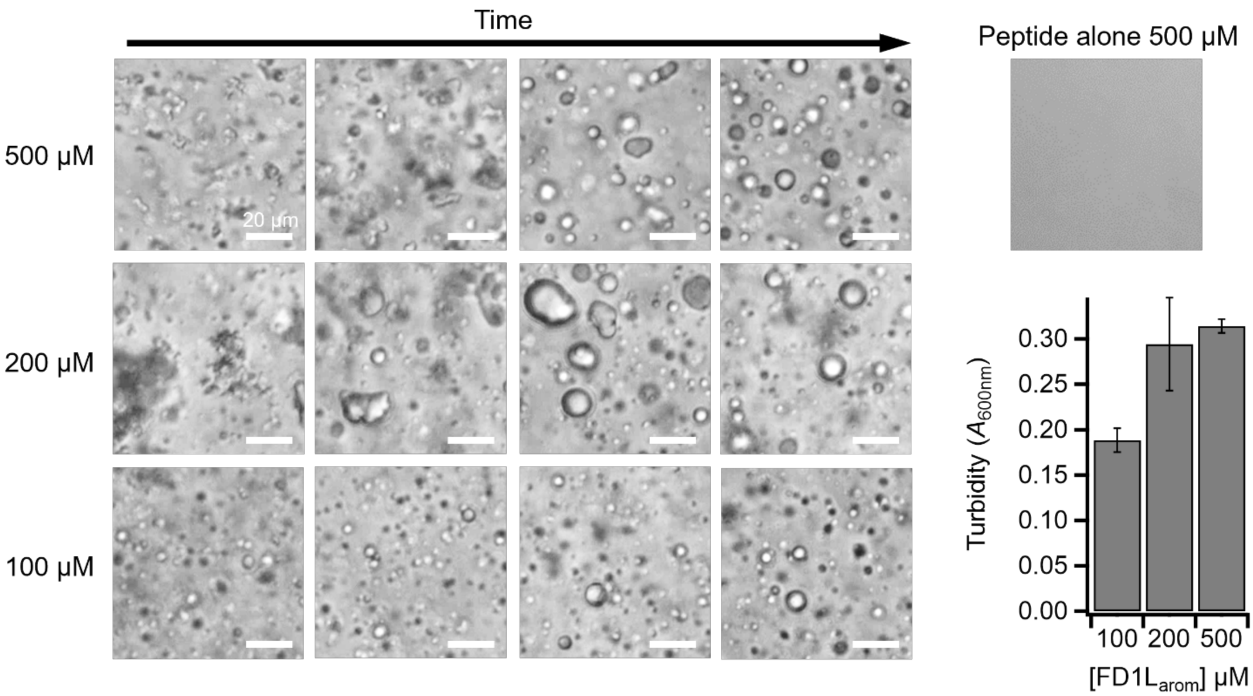


**Supplementary Fig. 5: Gel-like structures induced by peptides undergo a morphological transition from amorphous assemblies to spherical droplets upon temperature increase.** Representative DIC images of αSyn assemblies induced by FD1L$_{arom}$ are shown. The initial assemblies were prepared by incubating the samples on ice for 1 hour. Morphological changes were monitored over time at room temperature using DIC microscopy.

**a**

| Peptide | Sequence | Solubility score | Peptide | Sequence | Solubility score |
|---|---|---|---|---|---|
| FL2LΔ$_{posi2}$ | YKRWGKPRWWRRSYWS-NH$_2$ | 1.901 | FD1L$_{posi2}$ | YLGWHKLWFQPYRRRGGKKK-NH$_2$ | 1.796 |
| FL2LΔ$_{posi2,scr1}$ | SWRWYRWPRGKYKRSW-NH$_2$ | 1.962 | FD1L$_{posi2,scr1}$ | RGWKHRGLYLQYPFRWGKKK-NH$_2$ | 1.657 |
| FL2LΔ$_{posi2,scr2}$ | RGWPKSRSWKYYRWWR-NH$_2$ | 1.847 | FD1L$_{posi2,scr2}$ | HWPYKFQGRWLRLYRGGKKK-NH$_2$ | 2.116 |
| JMD$_{83-96}$ | KLKRKYWWKNLKMM-NH$_2$ | 2.142 | PolyK | YKKKKKKKKKKKKKKKKKKK-NH$_2$ | 5.242 |

**b**

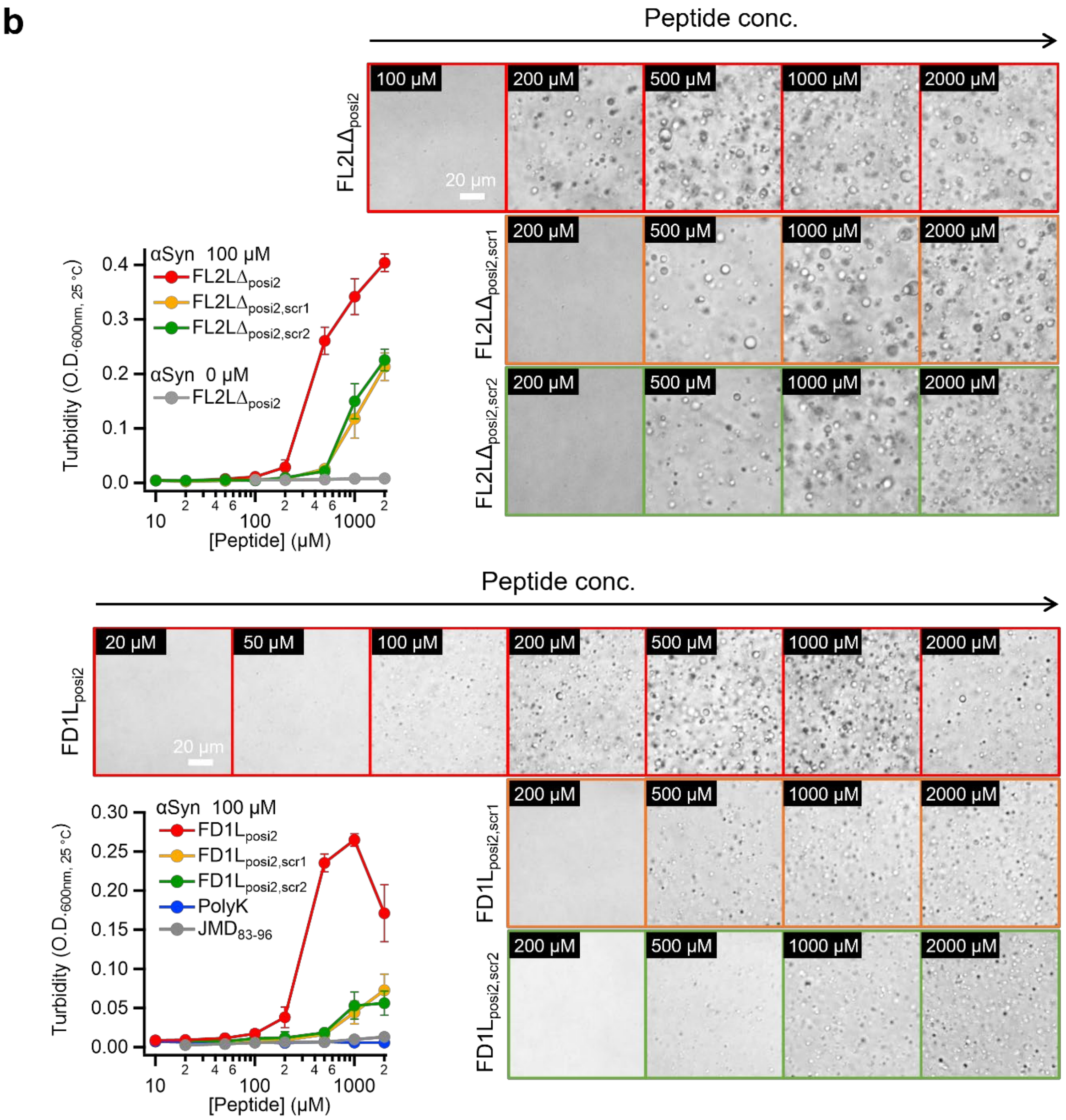


**Supplementary Fig. 6: Sequence scrambling of peptides significantly reduces LLPS efficiency.** **a** Amino acid sequence of scrambled variants of FL2LΔ$_{posi2}$ (FL2LΔ$_{posi2,scr1}$ and FL2LΔ$_{posi2,scr2}$) and FD1L$_{posi2}$ (FD1L$_{posi2,scr1}$ and FD1L$_{posi2,scr2}$). **b** Phase diagrams and representative DIC images of αSyn LLPS induced by the respective peptides, illustrating concentration-dependent phase behavior.

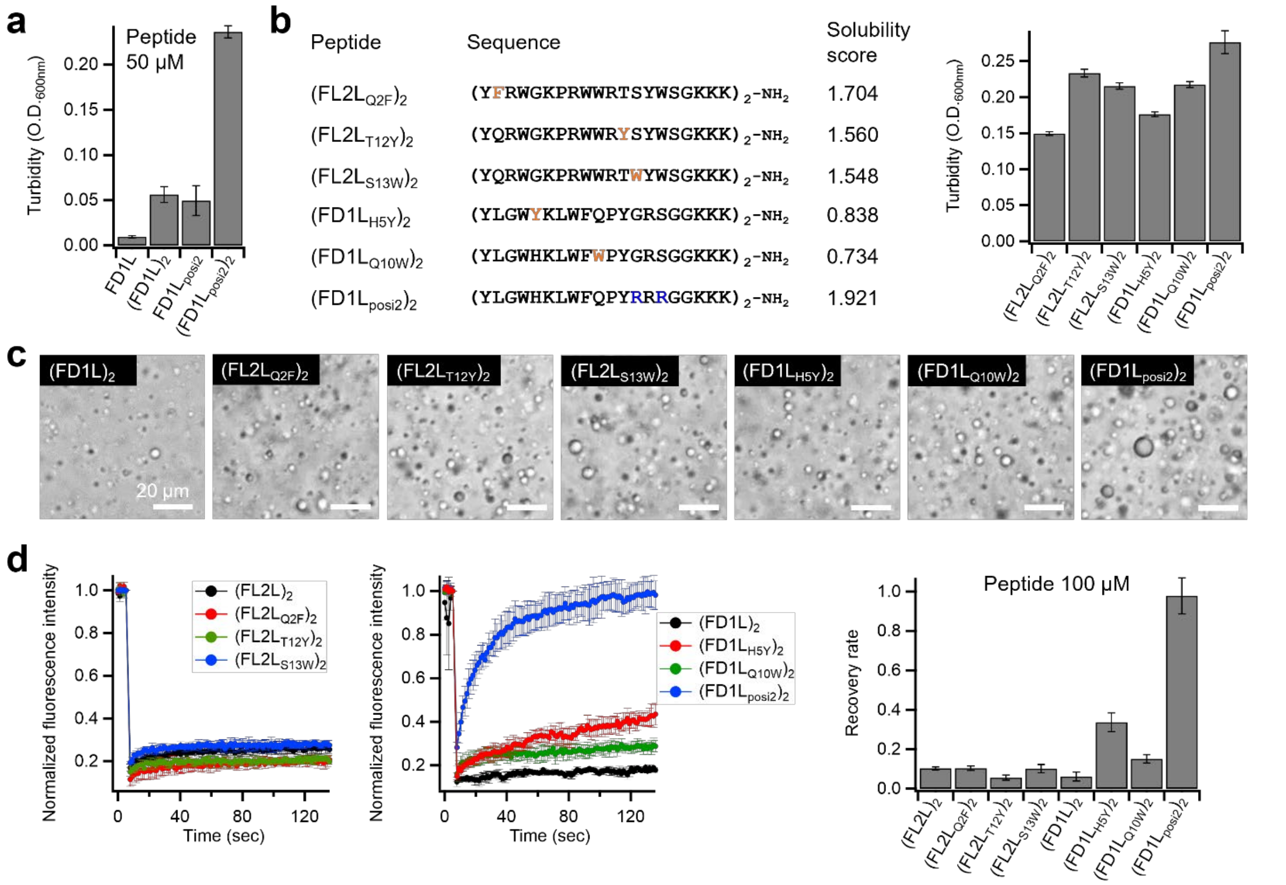

**Supplementary Fig. 7: Tandem repetition enhances LLPS efficiency while modulating droplet fluidity.**

**a** Comparison of turbidity in solutions containing 100 µM αSyn with 50 µM of either non-repetitive or tandem-repeat peptides: FD1L, (FD1L)$_2$ FD1L$_{posi2}$, (FD1L$_{posi2}$)$_2$. **b** Amino acid sequences of tandem-repeat peptide variants and their LLPS-inducing capacities evaluated via turbidity measurements. **c** Representative DIC images of αSyn droplets induced by tandem-repeat peptides. **d** FRAP recovery curves demonstrating that (FD1L$_{posi2}$)$_2$ maintains high droplet liquidity, whereas other variants exhibit reduced fluidity.

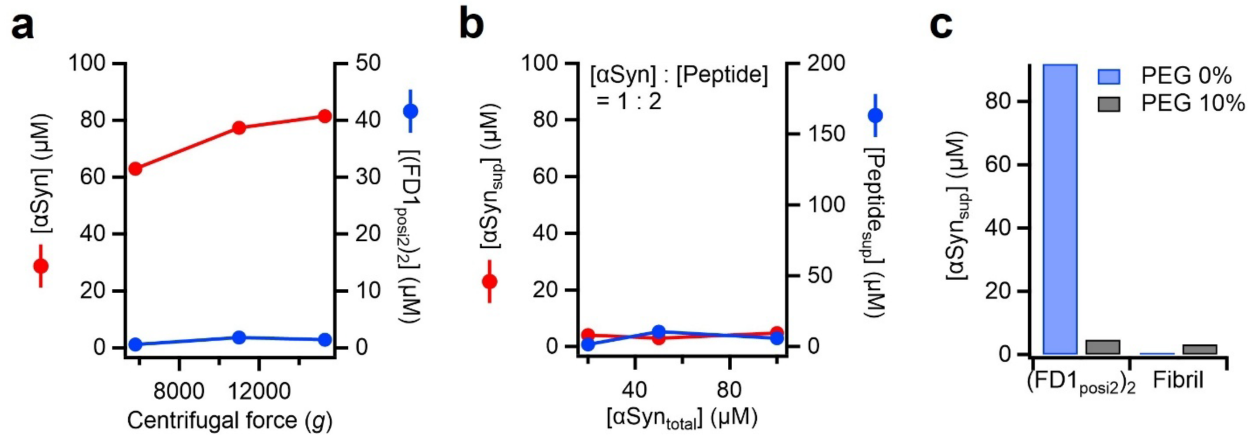


**Supplementary Fig. 8: Residual concentrations of αSyn and (FD1L$_{posi2}$)$_2$ in the supernatant after centrifugation.**

**a** Concentrations of αSyn and (FD1L$_{posi2}$)$_2$ remaining in the supernatant (αSyn$_{sup}$ and Peptide$_{sup}$, respectively) after centrifugation at 5,800×*g*, 11,000×*g*, and 15,300×*g*, determined by HPLC. **b** Supernatant concentrations of αSyn and (FD1L$_{posi2}$)$_2$ following centrifugation of solutions containing varying initial concentrations of αSyn and a two-fold molar excess of peptide. **c** Comparison of residual αSyn concentrations in the supernatant of solutions containing either LLPS-derived droplets or amyloid fibrils. The residual αSyn concentration was slightly higher in the droplet sample (4.65 μM) compared to the fibril sample (3.16 μM).

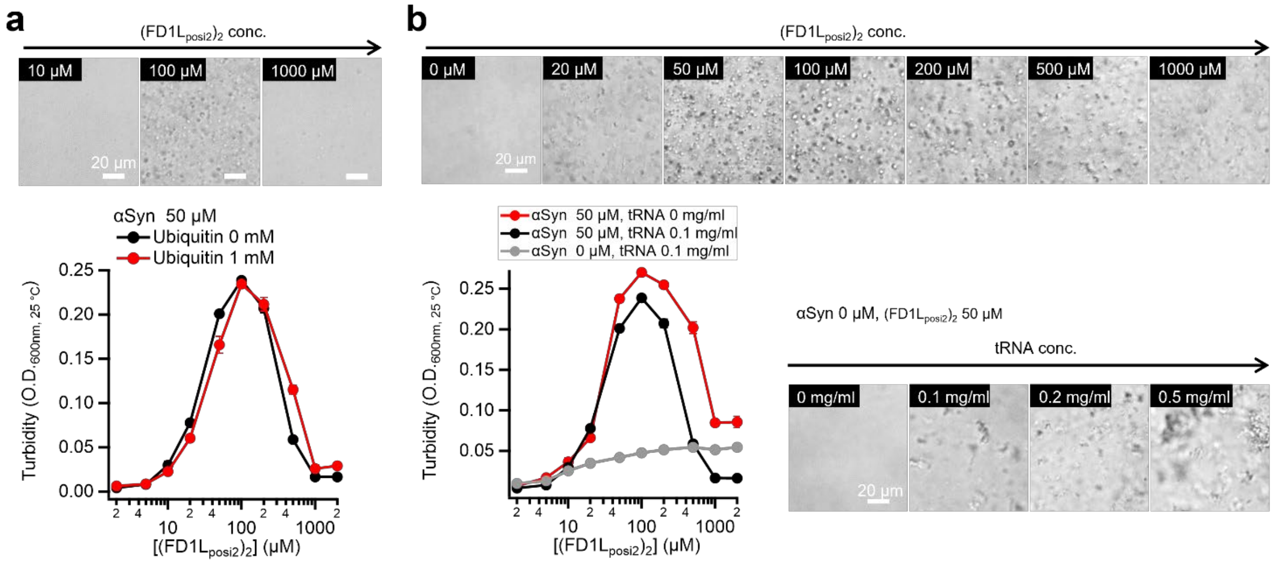


**Supplementary Fig. 9: Phase diagrams of peptide-mediated αSyn LLPS in the presence of other biomacromolecules.**

**a** Phase diagrams demonstrating that the presence of ubiquitin does not affect the LLPS behavior of αSyn. **b** Phase diagrams showing that tRNA does not alter the position of the LLPS peak; however, at high concentrations of $(FD1L_{posi2})_2$, the morphology of the condensates shifts toward an aggregate-like appearance. Notably, $(FD1L_{posi2})_2$ forms aggregates with tRNA even in the absence of αSyn.

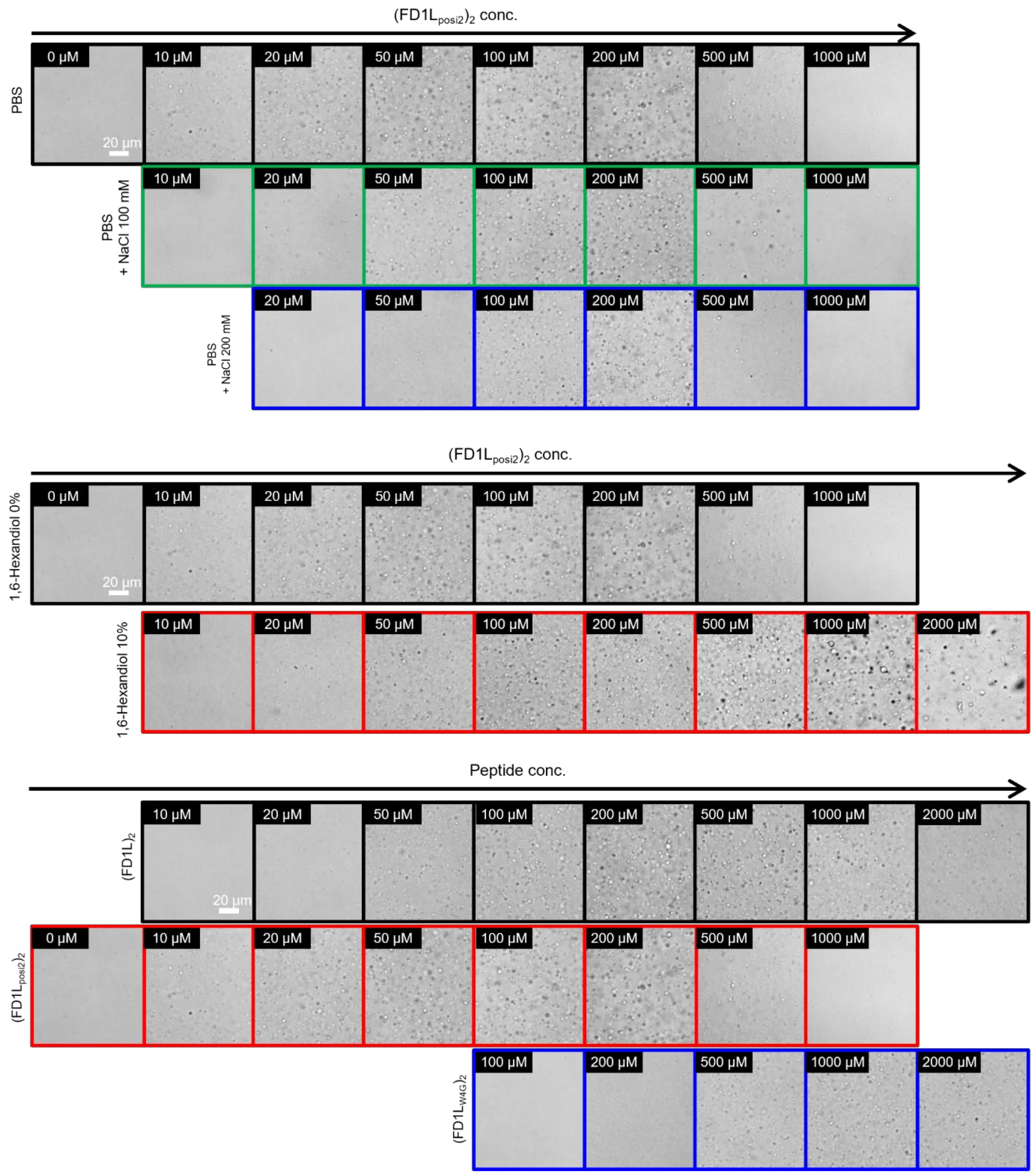


**Supplementary Fig. 10: Representative DIC images of solution prepared under various conditions as shown in Fig. 4c.**

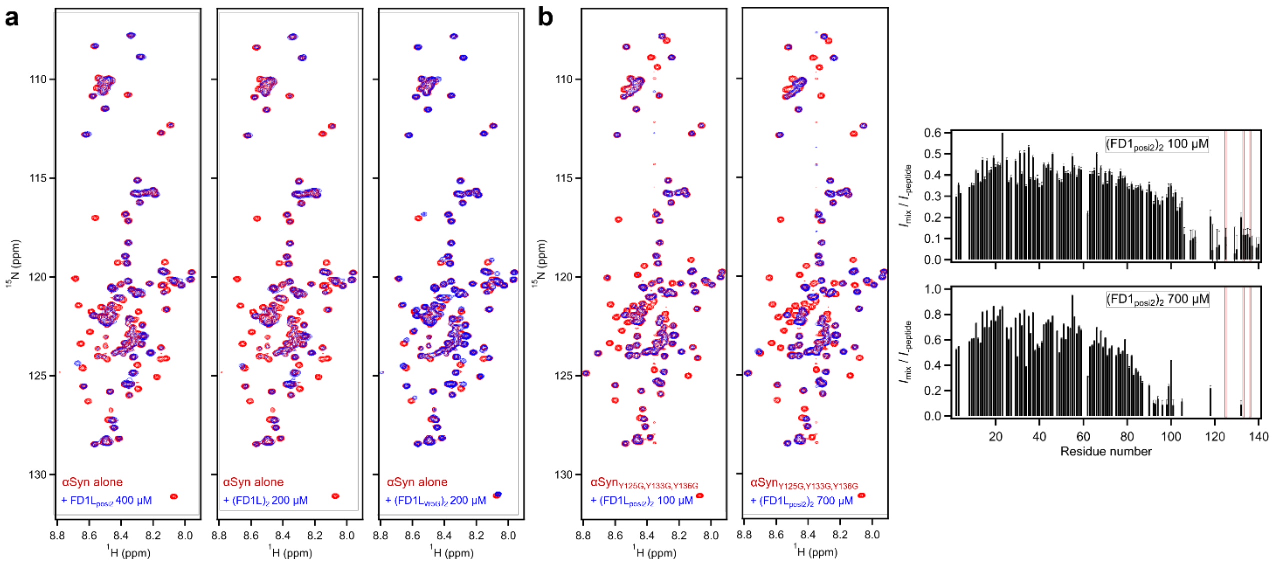


**Supplementary Fig. 11: HSQC spectra of αSyn in the presence of LLPS-inducing peptides.**

**a** Comparison of $^{1}$H–$^{15}$N HSQC spectra of αSyn recorded in the absence (red) and presence (blue) of FD1L$_{posi2}$, (FD1L)$_2$, and (FD1L$_{W5G}$)$_2$. Spectral changes highlight residue-specific perturbations associated with peptide binding. **b** Interaction between (FD1L$_{posi2}$)$_2$ and αSyn mutants in which Y125, Y133, and Y136 were replaced with glycine showed HSQC spectra largely similar to those of the wild-type αSyn, indicating that overall binding modes were not significantly altered. However, LLPS induction was markedly reduced in these mutants, demonstrating that the aromatic residues are critical for efficient condensate formation even though they contribute little to detectable chemical shift perturbations.

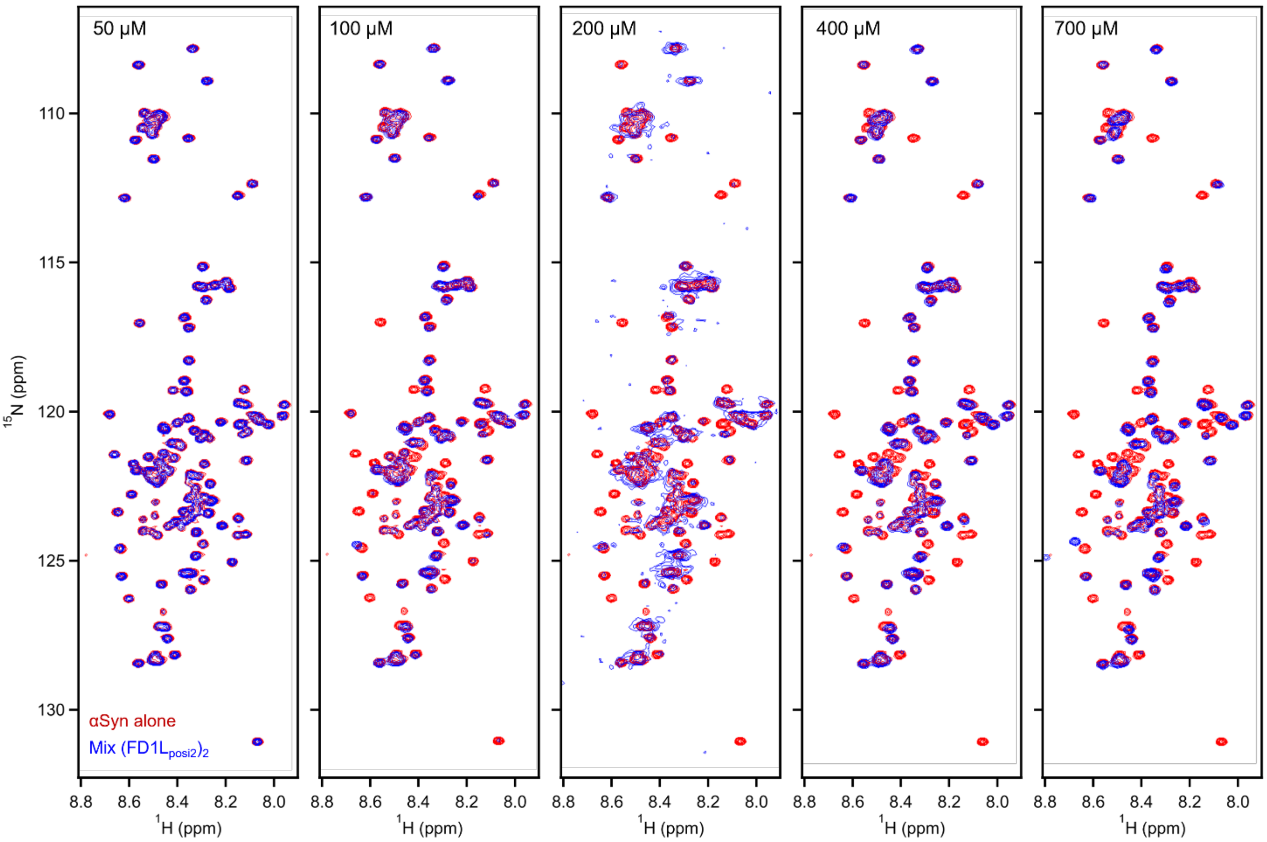


**Supplementary Fig. 12: HSQC spectra of αSyn at increasing concentrations of (FD1L$_{posi2}$)$_2$.** Progressive line broadening and intensity attenuation reflect concentration-dependent interactions and the onset of condensate formation.

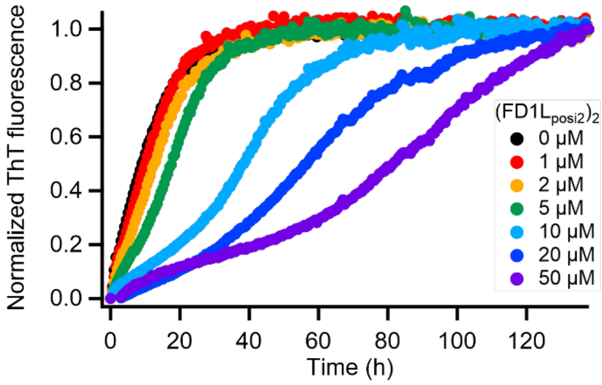


**Supplementary Fig. 13: Normalized ThT fluorescence profiles of αSyn fibril elongation in the presence of LLPS-inducing peptides.**

ThT fluorescence intensities from Fig. 6b were normalized to the endpoint signal to allow direct comparison of kinetic traces. The resulting profiles highlight the concentration-dependent inhibitory effect of peptides on fibril elongation, demonstrating that peptide binding slows the kinetics of amyloid formation.
